# Measuring integrin force loading rates using a two-step DNA tension sensor

**DOI:** 10.1101/2024.03.15.585042

**Authors:** J. Dale Combs, Alexander K. Foote, Hiroaki Ogasawara, Arventh Velusamy, Sk Aysha Rashid, Joseph Nicholas Mancuso, Khalid Salaita

**Author notes:** These two authors contributed equally to this work (J. Dale Combs, Alexander K. Foote).

## Abstract

Cells apply forces to extracellular matrix (ECM) ligands through transmembrane integrin receptors: an interaction which is intimately involved in cell motility, wound healing, cancer invasion and metastasis. These small (pN) forces exerted by cells have been studied by molecular tension fluorescence microscopy (MTFM), which utilizes a force-induced conformational change of a probe to detect mechanical events. MTFM has revealed the force magnitude for integrins receptors in a variety of cell models including primary cells. However, force dynamics and specifically the force loading rate (LR) have important implications in receptor signaling and adhesion formation and remain poorly characterized. Here, we develop a LR probe which is comprised of an engineered DNA structures that undergoes two mechanical transitions at distinct force thresholds: a low force threshold at 4.7 pN corresponding to hairpin unfolding and a high force threshold at 56 pN triggered through duplex shearing. These transitions yield distinct fluorescence signatures observed through single-molecule fluorescence microscopy in live-cells. Automated analysis of tens of thousands of events from 8 cells showed that the bond lifetime of integrins that engage their ligands and transmit a force >4.7 pN decays exponentially with a τ of 45.6 sec. A small subset of these events (<10%) mature in magnitude to >56pN with a median loading rate of 1.3 pNs^-1^ with these mechanical ramp events localizing at the periphery of the cell-substrate junction. Importantly, the LR probe design is modular and can be adapted to measure force ramp rates for a broad range of mechanoreceptors and cell models, thus aiding in the study of mechanotransduction.

**TOC:** 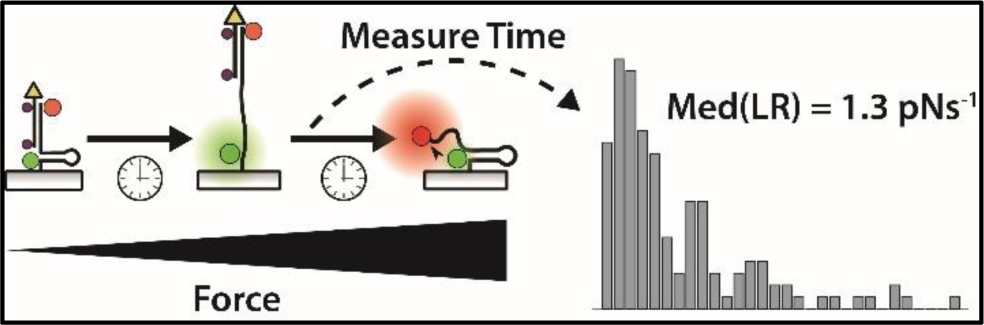

## Introduction

Many important biomolecular interactions involved in sustaining life are regulated through small, piconewton (pN) level forces. These forces, applied to and by the cells through membrane receptors such as integrins, play a pivotal role in transducing signals into downstream biochemical pathways, a process described as mechanotransduction.^1–4^ The process of interconverting mechanical cues into biochemical signals is critical to a vast array of biological processes, including wound healing, cell motility, and tissue fibrosis.^5–8^ Molecular tension fluorescence microscopy (MTFM) probes have been developed to measure and analyze the forces transmitted at cell-extracellular matrix (ECM) and cell-cell interfaces, employing extendable molecules such as PEG^9–10^, DNA^11^, and proteins^12^ that are flanked by fluorescent donor-acceptor pairs.^13–16^ Studies employing molecular tension probes have demonstrated that integrins sense and activate mechanotransduction pathway in response to pN forces.^17–21^ While existing probes have revealed the magnitude of forces transmitted by integrins, the force dynamics remain unclear, particularly at the molecular scale. Quantifying integrin force dynamics is critical because it is an important parameter in studying adhesions^22^ as well as in understanding how various integrin-targeting drugs modulate its function.^23–27^ Indeed, there has been much speculation about the integrin force loading rate (LR), which has been estimated from bulk measurements of the deformation of elastomeric substrates with values ranging from 0.007 pN/sec up to 4 pN/sec.^28^

One potential approach to measuring the LR is to perform time-dependent single-molecule measurements of MTFM probes. However, this is a challenge because it is difficult to characterize single-molecule fluorescence dynamics for weak and transient signals that are prone to photobleaching while in the presence of living cells that generate autofluorescence. To address these problems we created a force LR probe with two enabling features. First, the probe undergoes two digital mechanical transitions at distinct forces identified by unique fluorescence signatures (**Figure 1a**).^29–30^ This strategy avoids the ambiguity of analog sensors.^31–33^ Second, we engineered the LR probe with dual quenchers that significantly suppress photobleaching.

**Figure 1:**
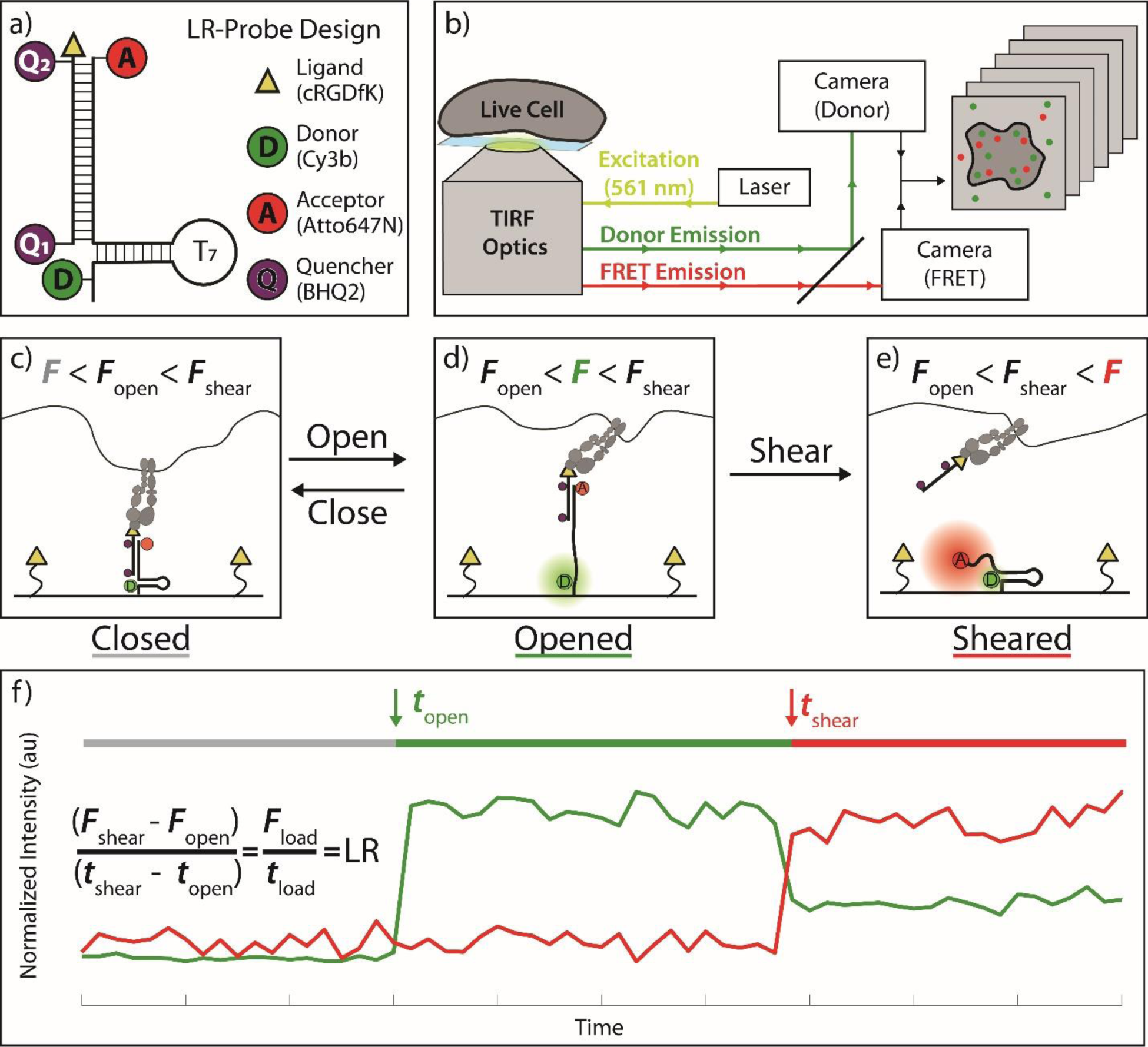
Schematic of LR probe and expected data. a) LR probe schematic. b) Optical diagram showing microscope setup used for single-molecule fluorescence microscopy of LR probe. c) Schematic showing initial integrin engagement with LR probe at low force (F<4.7 pN) resulting in the probe in the closed state. Both donor and FRET signals are quenched in the closed state. d) F> 4.7 pN but <56 pN place the probe in the opened state which is characterized by donor turn-on signal and lack of FRET signal. e) Forces > 56 pN result in probe shearing characterized by FRET emission and diminished donor signal. f) An idealized donor/FRET trajectory, showing times when the probe opened (*t*_open_) and sheared (*t*_shear_), and marking the opened and sheared states with a green and red line. Force values for opening and shearing and the timestamps of each transition are used to calculate the loading rate (LR).

The LR-probe was comprised of two oligonucleotide strands: a ligand strand and an anchor strand (**SI Note 1 – LR Probe Concept**, **SI Method 1 – LR Probe Synthesis**). The anchor folds into a hairpin and is modified with both a donor dye (Cy3B) and a FRET acceptor dye (Atto647N). The ligand strand presents a cyclic RGDfK (cRGD) peptide, contains two quenchers (Q_1_ and Q_2_), and is complementary to the anchor strand. The fluorescent output of the LR probe informs on its mechanical state and thus the force trajectory applied by the cell. Time-dependent single-molecule imaging of the LR-probe was performed by directly exciting the donor while simultaneously collecting fluorescence from both donor and FRET channels (**Figure 1b**, **Figure S1)**. When the LR-probe is in its closed state and little to no force is applied, both dyes are quenched reducing fluorescent output to near-background (**Figure 1c,f**). The hairpin domain of the LR-probe unfolds when *F* > 4.7 pN is applied, which is associated with early loading by integrin receptors (**Figure 1d**).^29^ Note that the 4.7 pN value represents the *F*_1/2_ which is the equilibrium unfolding force.^30^ This unfolding transition leads to a separation of the donor and Q_1_, resulting in a distinct turn-on signal of donor emission (**Figure 1d,f**). As the tension on the probe increases, the ligand strand undergoes irreversible shearing at *F* >56 pN which is the threshold for duplex shearing (**Figure 1e**).^34–35^ This second event leads to loss of the ligand strand containing both quenchers, allowing for probe refolding and a corresponding turn-on of FRET signal (**Figure 1e,f**). **Figure 1f** shows an idealized representation of fluorescence readout, and marks the timestamps of these transitions (*t*_open_ and *t*_shear_). By measuring the time between hairpin opening (*t*_open_, marked by donor turn-on) and duplex shearing (*t*_shear_, marked by FRET emission) using single-molecule fluorescence microscopy, we can estimate the loading rate between those two events in live cells within functional adhesions.

## Results

### Characterizing the LR probe

To validate the LR probe as single-molecule force sensor, we performed a series of controls to characterize the signal associated with its state transitions (closed, opened, and sheared, **Figure 1c-e**). The first control experiment characterized the signal associated with opening of the DNA hairpin with *F* > 4.7 pN (closed➜opened transition). This transition was mimicked by first measuring a closed LR probe on a surface, and then adding (at frame 0) a 17-mer complement (20 μM) to the hairpin stem-loop which irreversibly locks the DNA hairpin into the opened state. To anchor the LR probe to the glass surface, we grafted a dense layer of biotinylated-PEG onto the surface modified with streptavidin (SI Method 2 – LR Probe Surface Preparation).^36^ We subsequently added the LR probe (400 pM concentration) to achieve a sparse density of probes appropriate for single-molecule imaging (**Figure 2a**, **Figure S2, Figure S3**). **Figure 2a** shows representative fluorescent images from both the donor (green) and FRET (red) channels at time points which highlight the closed to opened state transition. These images show the entire field of view, with a white inset highlighting a 100×100 pixel region and a further yellow inset highlighting a 5×5 pixel region which marks the location of a single molecule. The donor and FRET signals within this 5×5 region were summed to show their intensity as a function of time (**Figure 2b**). From this trace, it is clear that the probe starts in the closed state (frames 0-13, low donor and FRET signal), transitions to an opened state (frames 14-60, high donor and low FRET signal), and exhibits single-step photobleaching of the donor (frames 61 onwards). Above the intensity trajectory in **Figure 2b** we mark with a green line the opened state probe which is followed by donor photobleaching. Further controls were performed quantifying the negligible bleed-through of the donor dye into the FRET channel (**Figure S4**), the rate of donor photobleaching (**Figure S5**), and the kinetics of the LR probe opening through its complement locking strand (**Figure S6**). The kinetics of hairpin locking are similar to those previously characterized in ensemble measurements.^37^

**Figure 2:**
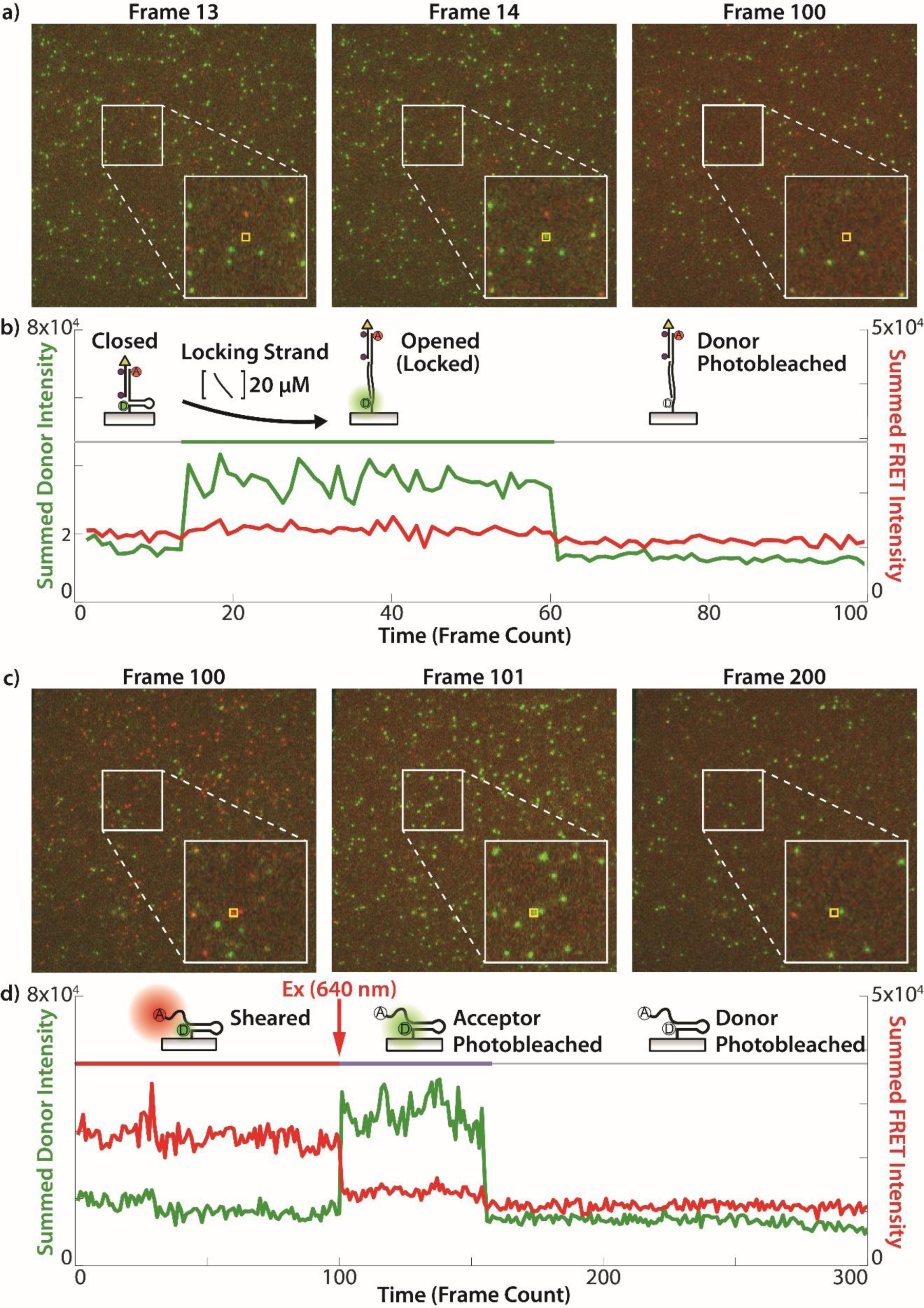
LR probe single-molecule fluorescence emission validation. a) Representative images from a timelapse of a surface with LR probe showing donor (green) and FRET (red) fluorescent signal. White inset shows a 100×100 pixel region, and the further yellow inset shows a 5×5 region highlighting a single molecule. Frame 13 shows the probe in the closed state, Frame 14 the probe is locked into the opened state with its complement strand, and Frame 100 the signal has photobleached. b) Time trace of the fluorescent signal within the yellow 5×5 region marked in 2a, with the probe’s state assignment shown with the horizontal colored bar. c) Representative images from a timelapse of a surface with a sheared LR probe which begins in a high-FRET state with insets matching those in 2a. Frame 100 shows the probe in a high-FRET state, photobleaching is induced between Frame 100 and 101, and Frame 101 shows diminished FRET and recovered donor signal. By Frame 200 the donor has photobleached. d) Time trace of the fluorescent signal from the experiment shown in 2c.

Having identified the signal produced by both the closed and opened states of the probe, we next characterized the signal produced by the sheared state which occurs at *F* >56 pN (opened➜sheared transition). Shearing of the ligand strand allows the DNA hairpin to refold without the dual quenchers. This process activates the FRET between Cy3B and A647N, turning-on the FRET signal while diminishing the Cy3B emission. The sheared state of the LR probe was formed by attaching the probe to the surface without the top ligand strand (**Figure 2d**, probe drawing). **Figure 2c** shows representative images of the fluorescent readout, with white and yellow insets marking the corresponding 100×100 and 5×5 pixel regions as described previously. The fluorescence image at frame 100 shows many probes with the majority displaying high FRET, indicative of the sheared state. To induce photobleaching of the acceptor, we excited the sample with 640 nm high-intensity illumination between frames 100 and 101. This photobleaching of the acceptor shows conversion of FRET puncta into donor emission puncta, further confirming that the signal is due to single-molecule LR probes. This conversion can be visualized by comparing frames 100 to 101 in **Figure 2c**. As before, a representative timetrace of donor and FRET intensities are shown in **Figure 2d** with the probe beginning in the sheared state (frames 0-100, low donor and high FRET). Next, acceptor photobleaching occurs after frame 100, diminishing FRET signal and causing an increase in donor Cy3B emission (frames 100-155, high donor and low FRET). Finally, the Cy3B donor photobleaches, leading to single-step termination of the donor signal (frames 156-onwards, low donor and low FRET). Above the intensity trajectory in **Figure 2d** we mark with a red line the sheared state, and we mark with a light-purple line the acceptor photobleached state. Importantly, the low donor/high FRET signal characterizing the sheared state (**Figure 2d**, red line) is starkly different from the high donor/low FRET signal representing the opened state (**Figure 2b**, green line). Lastly, all traces show single-step transitions between states, and single-step termination due to photobleaching, validating that these characteristic signals are identifiable at the single-molecule level. Further controls quantified the FRET efficiency of the sheared state, yielding FE=85.5%+/-7% which is consistent with the donor-acceptor distance of the sheared state (**Figure S5**). Finally, a simulation using oxDNA was run, determining a (*F*_shear_ – *F*_open_) force-difference of 47.7 pN, comparable to the expected 51.3 pN for the LR probe (**Figure S7**, **SI Method 3 – OxDNA modeling of the LR Probe**).

### Single-molecule LR probe measurements in cell adhesions

We next performed experiments measuring live-cell integrin forces on the LR probes. We used NIH-3T3 fibroblast cells for our live cell experiments as fibroblasts are one of the most extensively studied models for integrin mechanotransduction.^38^ Cells were seeded onto the LR probe surface for 25 min (37C, 5% CO_2_, and 5% FBS FluoroBrite DMEM) allowing cells to spread then timelapse TIRF imaging was initiated. We alternated between RICM imaging and simultaneous donor/FRET imaging at a rate of 0.1 Hz over the course of the experiment spanning up to 100 min (**Figure 3a**). **Figure 3b** shows an overlay of the three channels at the initiation of imaging (*t*=0 min). Since the cells had already begun interacting with the surface, fluorescent signal from previously opened/sheared probes can be seen, with spatially distinct, puncta indicating LR-probe detection under single-molecule conditions (**Figure 3b**, **Figure S8**). To study the loading rate exerted by NIH-3T3 cells we identified single-molecule traces which followed the specific sequence of transitions described above (**Figure 1f**): closed state➜ opened state (4.7 pN) ➜ sheared state (56 pN). These transitions were detected by distinct fluorescence-signal outputs: a dark state (marking the closed state), an increase in Cy3B donor signal (marking the closed-to-opened state), and finally a simultaneous decrease in Cy3B donor signal and increase in FRET signal (marking the opened-to-sheared transition). A representative trace is shown in **Figure 3c**, with its spatial position relative to the cell marked by an arrow in **Figure 3b**. Additionally, **Figure 3c** marks the opened, sheared, and closed states with a green, red, and gray line, respectively, above the intensity trace timestamp. These transitions and their timestamps are amenable to manual quantification; however, we created custom code to fully automate analysis and increase the throughput to handle large data sets containing tens of thousands of puncta across hundreds of frames.

**Figure 3:**
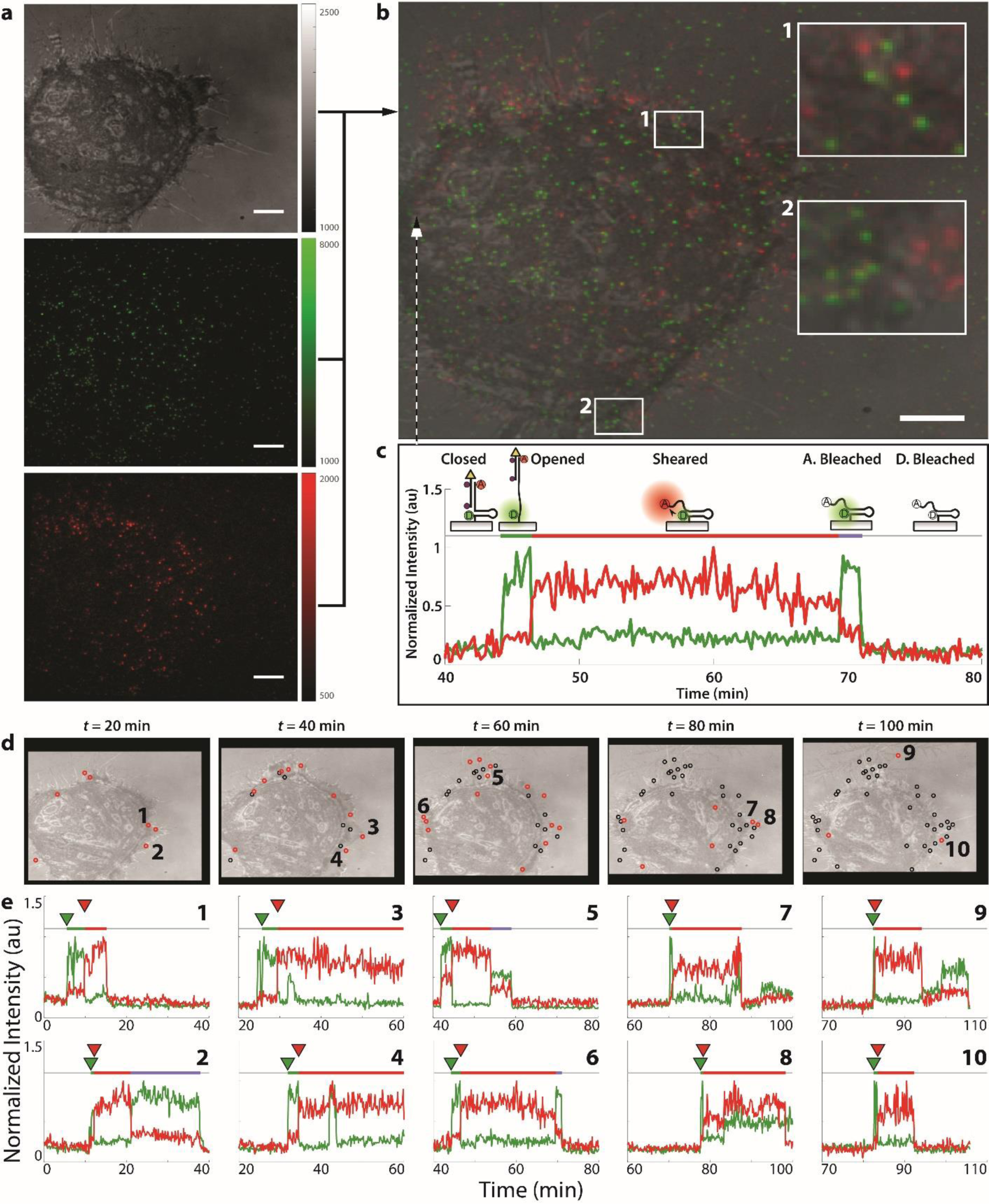
Single-molecule analysis of live-cell adhesions using LR probe. a) RICM (top) donor (middle) and FRET (bottom) images at *t*=0. Scale bar = XX micron, and the vertical intensity bars show raw EMCCD pixel intensity. b) Overlay of images from 3a, with two highlighted regions showing single-molecule puncta observed. c) Fluorescence time-trace of the 3×3 pixel region shown with an arrow. This time-trace exhibits the closed➜opened➜sheared transitions as seen in the donor (green) and FRET (red) intensities, and marks the opened, sheared, and acceptor photobleached state of the probe with a green, red, and purple horizontal line, respectively. d) Images of the cell at 20-minute intervals, with red circles marking locations of shearing events occurring during that interval. Shearing events are carried over between frames as black circles showing a buildup of events. Numbers 1-10 correspond to the fluorescent trajectories shown below. e) Fluorescent trajectories matching the color schematic in 3c, and with green and red arrows showing the timestamps of the opening transition (*t*_open_) and shearing transition (*t*_shear_), respectively. Locations are marked in Figure 3d.

Our custom Matlab codes are extensively described in **SI Note 3 – Automated Analysis of Single-Molecule Data**. Briefly, each point of signal was fit to a 2D-Gaussian to find spatially distinct single-molecule events. Next, intensities belonging to individual LR probes were identified and clustered together, and time traces for these single-molecule events were analyzed with a changepoint algorithm to identify state transitions and their kinetics.^39^ The evolution of the cell’s interaction with the surface, and the localization of real-time shearing events of LR probes is visualized in **Figure 3d**. Each subsequent image in **Figure 3d** shows an overlay of three channels (RICM, donor, FRET) at 20-minute intervals, and marks the location of a detected shearing event with a red circle. Shearing events are carried over between frames as black circles, so the history of high-force interactions can be visualized at later timeframes. This sequence of images has been reproduced into a video with the full time-resolution (**SI Video 1**), along with each single-molecule fluorescence trace (**SI Video 2**). We recommend that readers watch these videos now if able, as they helps visualize the raw data while also displaying the wealth of information made available through single-molecule experiments using the LR probe. As can be seen in **SI Videos 1 and 2,** and **Figure 3d**, the vast majority of shearing events (*F* > 56 pN) occur near the periphery of the cell, a phenomenon repeatedly observed in past work.^17, 29, 40^ In addition to the spatial information obtained from identifying shearing events, temporal information on the cells force loading rate can be derived from the fluorescence intensity traces. Ten representative traces exhibiting a shearing event are shown in **Figure 3e**, with a number 1-10 indexing their spatial coordinates according to **Figure 3d** above. Within the traces, a green arrow indicates the time of probe opening (*t*_open_) and the red arrow indicates the time of probe shearing (*t*_shear_) and these are automatically derived from the changepoint analysis. The color of the horizontal bar above the plot indicates the state of the probe: a green line representing the opened state, a red line representing the sheared state, and a light-purple line (when applicable) representing acceptor photobleached state before donor photobleaching (which terminates the signal). Importantly, the duration of the opened state (length of green line and the time difference between the green and red arrows, *t*_shear_-*t*_open_) is used to quantify the loading time, and hence the loading rate.

By repeating the analysis shown in **Figure 3**, we identified 138 traces from 8 NIH-3T3 cell time lapses which displayed the opening➜shearing transition and quantified their loading times (**Figure 4a**). Observed loading times ranged from 1-30 frames, with mean, median, and modal loading times of 66, 40, and 20 seconds, respectively. In all cases the loading rates were shorter than the average photobleaching rate of Cy3B under our conditions (71 frames, **Figure S5**). To derive the LR we assume a linear loading rate between the initial hairpin opening (4.7 pN) and duplex shearing (56 pN), and divide this force difference by the observed loading time (**Figure 4a**, equation and top x-axis). This calculation results in an observed median loading rate of 1.3 pNs^-1^ (with interquartile range of 0.62 – 2.80 pNs^-1^) exerted by live cell integrins under initial adhesion formations.

**Figure 4:**
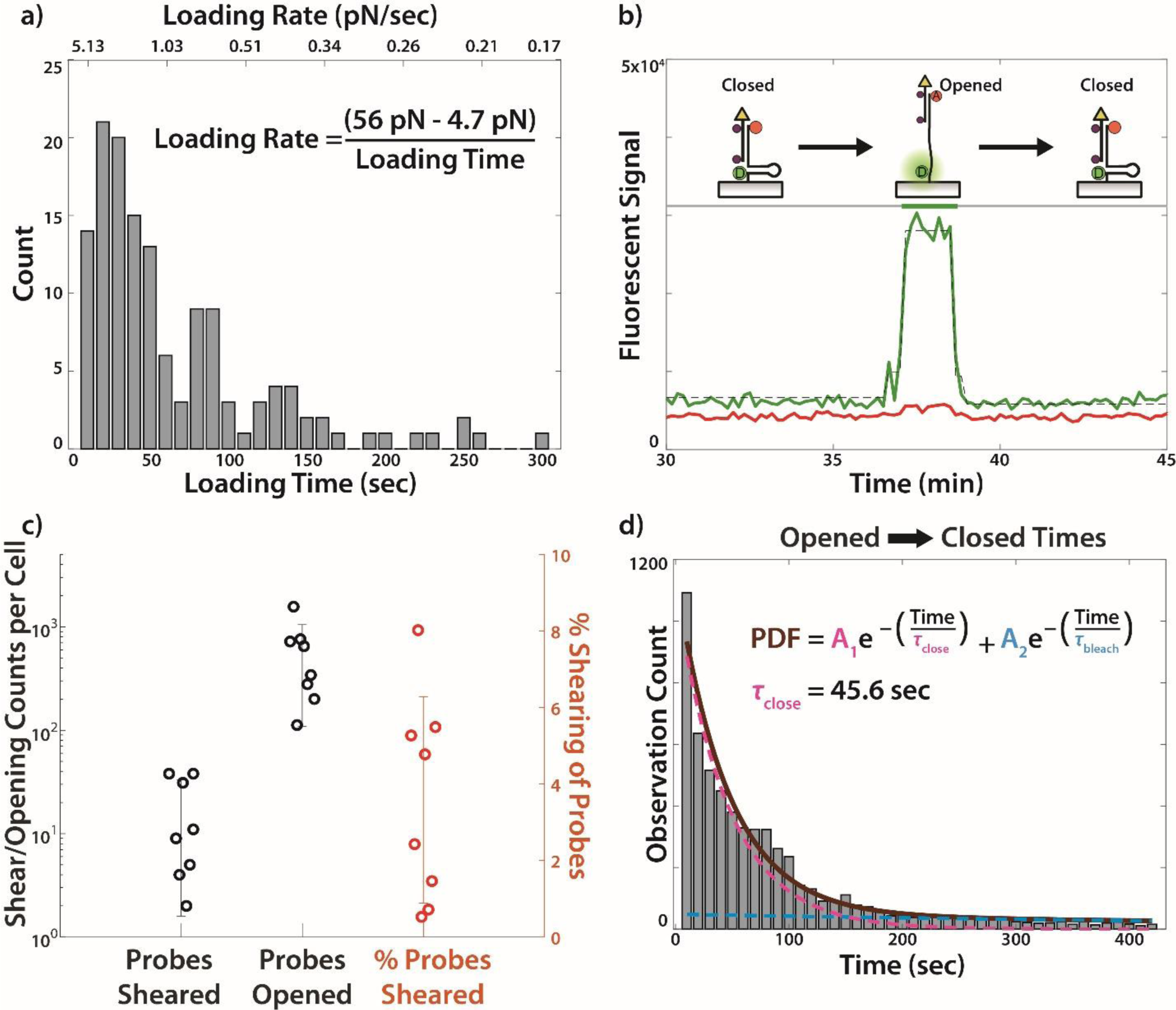
Analysis of LR probe live cell dynamics. a) Histogram of loading times (bottom axis). Inset equation shows how these values are converted to a loading rate (top axis). b) Fluorescent time-trace of a closed➜opened➜closed event, with a depiction of the probe states. Donor fluorescence is in green, FRET in red, and a dotted black line marks the changepoint algorithm used to detect the transition. This trace was obtained from the timelapse shown in Figure 3. c) Plot depicting LR probe event counts from 8 cells imaged for a duration of 100 min or less. The plot shows shearing and opening events for each cell as well as the ratio of opening events that lead to shearing events. d) Histogram of times for each opened event (N = 5,981) obtained from 8 cells. The distribution was fit to a biexponential (solid brown) composed of a fixed photobleaching rate (dotted blue) and variable closing time (dotted pink), where the weights of both exponentials (A_1_, A_2_) were allowed to vary.

While the focus thus far has been on the closed➜opened➜sheared transitions as quantified by the LR probe, the measurements also contain information on the closed➜opened➜closed transitions (**Figure 4b**). These events represent forces exerted by the cell which exceed 4.7 pN, but never reach the 56 pN threshold required to shear the probe. Identifying these events (See **SI Note 3.5 – Identifying Closed**➜**Opened**➜**Closed Transitions**) allows us to compare the number of “probes sheared” events to the number of “probes opened” events; a proxy to the dynamics of integrin adhesion formation which are often described by binding rates **(Figure 4c**).^41–42^ On average 3.6% of opened probes resulted in a shearing event, with some highly active cells shearing as many as 8.0% of opened probes (**Figure 4c**). In addition to the number of “probes opened” events observed, we also quantified the time during which the probe remained open before transitioning back to a closed state (as represented in **Figure 4b**). The timescales for these closed➜opened➜closed events are histogrammed in **Figure 4d** (N = 5981), and the probability distribution function (PDF) of all events < 420 sec were fit to a bi-exponential decay (brown line) composed of the rate of photobleaching (blue dotted line, *τ*_bleach_ = 710 sec, **Figure S5**) and the rate of closing (pink dotted line, *τ*_close_ = 45.6 ± 7.0 sec, 95% CI), with relative amplitudes of A_1_ and A_2_, respectively. A video showing the spatial location of closed➜opened➜closed transitions is included in **SI Video 3**, and demonstrates that most events are at the cell periphery, consistent with the regions of greatest mechanical activity.^17, 29^

## Discussion

The loading rate of integrin traction forces is an important metric in describing and predicting focal adhesion dynamics^25^. Therefore, we anticipate that LR measurements may improve integration between experiments and molecular clutch modeling of adhesions. The LR probe reveals several features of integrin force dynamics. First, only a small subset of integrin that initially engage RGD ligands and apply low levels of force (>5 pN) proceed to ramping force to 56 pN. Over 90% of integrin-ligand complexes that experience F>5 pN return to the idle state at low force. Second, the lifetime of the low-magnitude mechanical force (>5 pN) decays exponentially and displays a characteristic lifetime of 45.6 sec. This is consistent with integrin-ligand lifetime measurements that have ranged from 1-100s of seconds^43–45^. We note a cluster of force lifetimes between 80 and 100 secs that are more frequently observed than would be anticipated from the exponential decay function in **Figure 4d** and this may due to a long-lived integrin state^43^. Force may terminate due to simple ligand-receptor dissociation, as the k_off_ rate of integrin-RGD bonds are well documented^46^. Alternatively, the hairpin may refold due to termination of the force for ligated integrins. The highly dynamics mechanical nature of integrin-ligand forces reported here is consistent with reports of integrin lateral mobility and turnover within focal adhesions^47–49^. Third, we find that force loading to 56 pN is primarily localized to the cell periphery which spatially colocalizes with the area that displays the greatest frequency of hairpin opening events. Finally, force LR is highly heterogeneous with a distribution about a median of 1.3 pNs^-1^. This LR is consistent with reports by Sheetz and Roca-Cusachs that estimated a loading rate of 2.5 pNs^-1^ by inference of actin retrograde flow measurements in conjunction with measurements of the force required to unfold talin^24^. The same group also used loading rates obtained from traction force microscopy to infer individual integrin loading rates of 0.007 to 4 pN/s.^28^ Note that this estimate is derived from time-dependent stress measurements and requires an assumption of the density of integrins which can range from tens to hundreds per square micron.

Past work with single-molecule force spectroscopy, such as optical tweezers and AFM, apply force to integrin-ligand bonds and record the bond duration leading up to rupture. These values provide insights into bond lifetime under force, rather than the actual force lifetime or the native loading rate of integrins-ligand complexes within adhesions. Importantly, our LR Probe measures forces generated by the cell itself and transmitted to single integrins, and the observed force loading rates are orders of magnitude lower in magnitude compared to those induced through AFM and optical/magnetic bead techniques ^28, 50–51^. Given that ligand-receptor rupture force is highly dependent on loading rate, our work suggests that much of the single-molecule force spectroscopy data needs to be adjusted for physiological loading rates.

Challenges inherent to single-molecule measurements warrant discussion. First, to accommodate the long timescale of the experiment (> 1hr), the framerate was set to 0.1 Hz, or every 10 seconds. Any events undergoing their transition sequence (e.g. closed➜opened➜sheared) faster than our framerate would remain unobserved. From **Figure 4a** it is clear that the bulk of the loading time distribution is successfully measured, but care must be taken when selecting an acquisition rate, and should be optimized based on the observed rates of a given system. Second, a large number single-molecule events (fluorescent puncta) are observed which do not follow the described sequence transition (closed➜opened➜sheared). Many photophysical or biological events can alter the fluorescent output at the single-molecule level, such as photobleaching, dye blinking, probe degradation, heterogeneous microenvironments (including the nuclease rich environment under the cell), etc. While it is certainly feasible to quantify all observed single-molecules and ascribe their fluorescent trajectories to a given interference, we opted to utilize the distinct output of the LR probe to assign a strict selection criteria. In other words, the analysis algorithm was set for high specificity to select closed➜opened➜shearing events, and this high fidelity comes at the cost of ignoring many events, some of which may be true shearing events. Lastly, we note one possible false shearing signal in the scenario of Q1 bleaching, while Q2, donor and acceptor remaining active. This would appear as a long-lived opening event that could progress to a shearing, if such an event occurred rapidly after the initial bleaching, and would hence provide an inflated loading time (low LR). However, such a sequence of events occurring on the same probe is highly unlikely under the given experimental parameters.

## Conclusion

In summary, we have developed a novel method for detecting two distinct transitions of a single molecular complex comprised of two DNA strands by fluorescence de-quenching followed by FRET. This probe can detect both low (>4.7 pN) and high (>56 pN) force thresholds which occur during the same integrin-receptor interaction at the single-molecule level. We derive loading rate measurements for the live cell integrin mechanoreceptors which are congruent with estimates modeled in the field. This methodology protects both fluorophores from photobleaching and enables high fidelity detection of sequential mechanical transitions of DNA strands of single molecular complexes. The LR probe can be adapted to detect different thresholds by engineering the nucleic acid geometry or DNA sequence and such tuning is needed to investigate different classes of mechanoreceptor. Finally, we note that the LR probe is not specific to integrins and the ligand can be replaced with a wide variety of peptides or proteins appropriate for the biological system of interest.

## Supporting information

SI Video 1: Localizations of shearing events marked with red circles overlaid with an RICM image of cell 1

SI Video 2: Timelapse including SM fluorescent traces

SI Video 3: Localization of all closed->opened->closed events

SI Video 4: RICM Cell 2

SI Video 5: RICM Cell 3

SI Video 6: RICM Cell 4

SI Video 7: RICM Cell 5

SI Video 8: RICM Cell 6

SI Video 9: RICM Cell 7

SI Video 10: RICM Cell 8

## ASSOCIATED CONTENT

The following files are available free of charge here: https://www.dropbox.com/scl/fo/wcesxysdg3uigtwajz57a/h?rlkey=l9swiyeeersoxuvjz50barnec&dl=0

### Supplementary Information (.pdf)

**Supplementary Video 1** (.avi): Localizations of shearing events marked with red circles overlaid with an RICM image of cell #1.

**Supplementary Video 2** (.avi): The same RICM timelapse as in Supplementary Video 1 (left) but with the normalized fluorescence intensity time traces of donor (green) and acceptor (red). Vertical green and blue lines mark the timestamps of t_open_ and t_shear_, respectively. The localization of the single-molecule time trace (right) is shown as a blue circle on top of the RICM (left).

**Supplementary Video 3** (.avi): RICM timelapse of cell #1, with red circles marking locations of closed➜opened➜closed transitions as they are identified through time.

**Supplementary Video 4** (.avi): Localizations of shearing events marked with red circles overlaid with an RICM image of cell #2.

**Supplementary Video 5** (.avi): Localizations of shearing events marked with red circles overlaid with an RICM image of cell #3.

**Supplementary Video 6** (.avi): Localizations of shearing events marked with red circles overlaid with an RICM image of cell #4.

**Supplementary Video 7** (.avi): Localizations of shearing events marked with red circles overlaid with an RICM image of cell #5.

**Supplementary Video 8** (.avi): Localizations of shearing events marked with red circles overlaid with an RICM image of cell #6. **Supplementary Video 9** (.avi): Localizations of shearing events marked with red circles overlaid with an RICM image of cell #7.

**Supplementary Video 10** (.avi): Localizations of shearing events marked with red circles overlaid with an RICM image of cell #8.

## AUTHOR CONTRIBUTIONS

The manuscript was written through contributions of all authors. All authors have given approval to the final version of the manuscript. ‡These authors contributed equally.

## ACKNOWLEDGMENTS

K.S. acknowledges support from NIH NIAID R01AI172452 and NIGMS R01GM131099 and 1RM1GM145394.

