## Supplementary Figures and Text for "Measuring integrin force loading rates using a two-step DNA tension sensor"

‡These authors contributed equally.

**SI Figures**

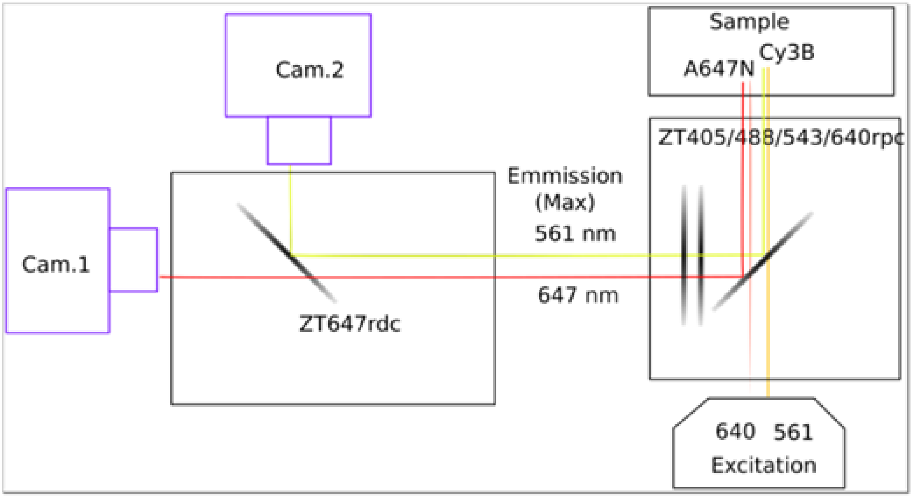


**Figure S1.** Optical configuration for emission resolved FRET. Laser lines of 561 and 647 nm are indicated as well as dichroic mirrors, emission filters, sample, and cameras to image Cy3B emission (Cam. 2) and A647N emission (Cam. 1).

**These files are being withheld by the authors until after peer-reviewed publication.**

**Figure S2.**

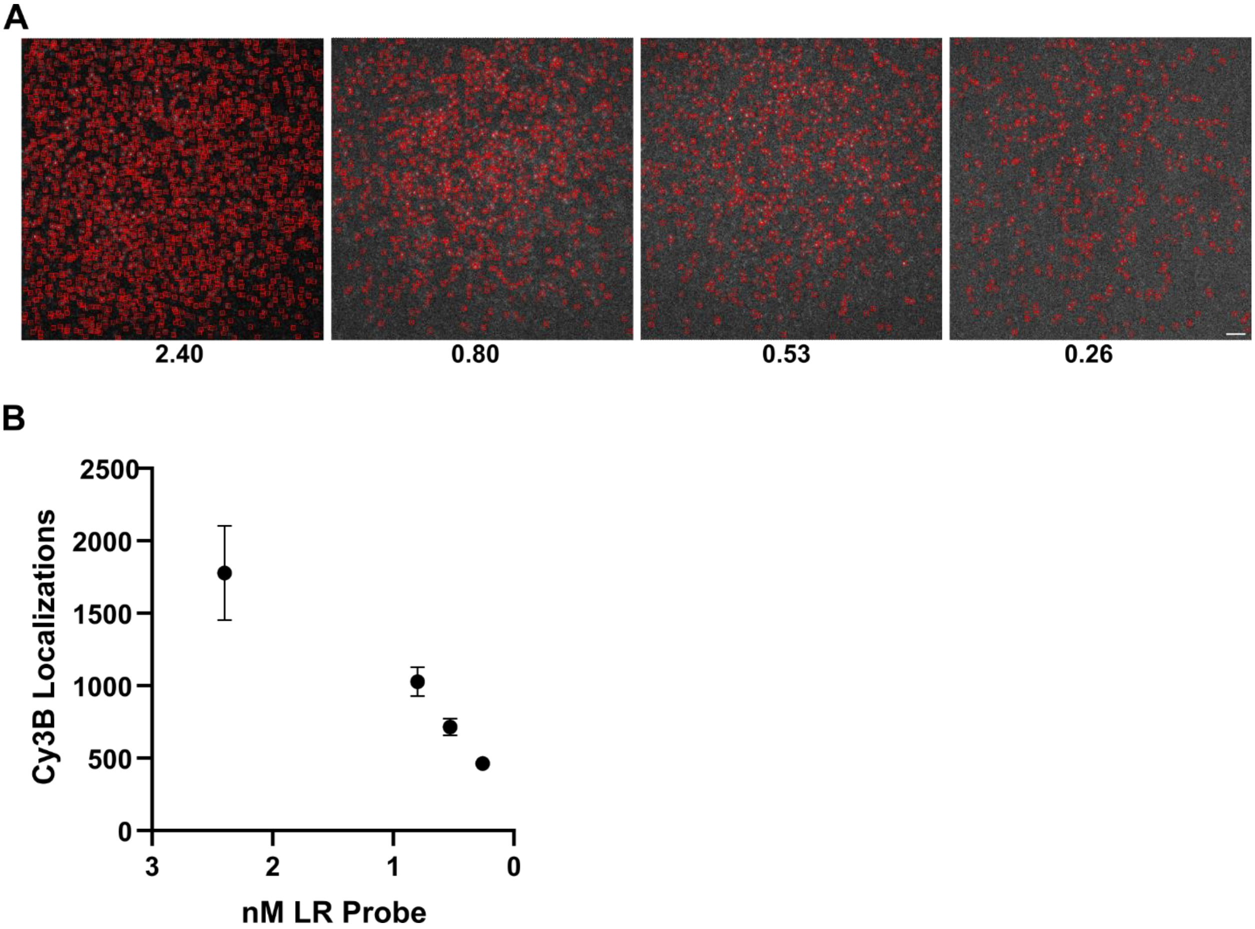


**Figure S3.** Titration of biotinylated surfaces with different concentration of LR probe. (A) Representative micrographs of surfaces exposed to 2.40, 0.80, 0.53, and 0.26 nM hybridized ligand and anchor strand LR probe with localizations boxed in red by Picasso software using settings of box Size = 7, Min. Net Gradient: 19700.[1] (B) Plot of localizations vs concentration of LR probe exposure for three Cy3B fluorescence micrographs of one surface per concentration.

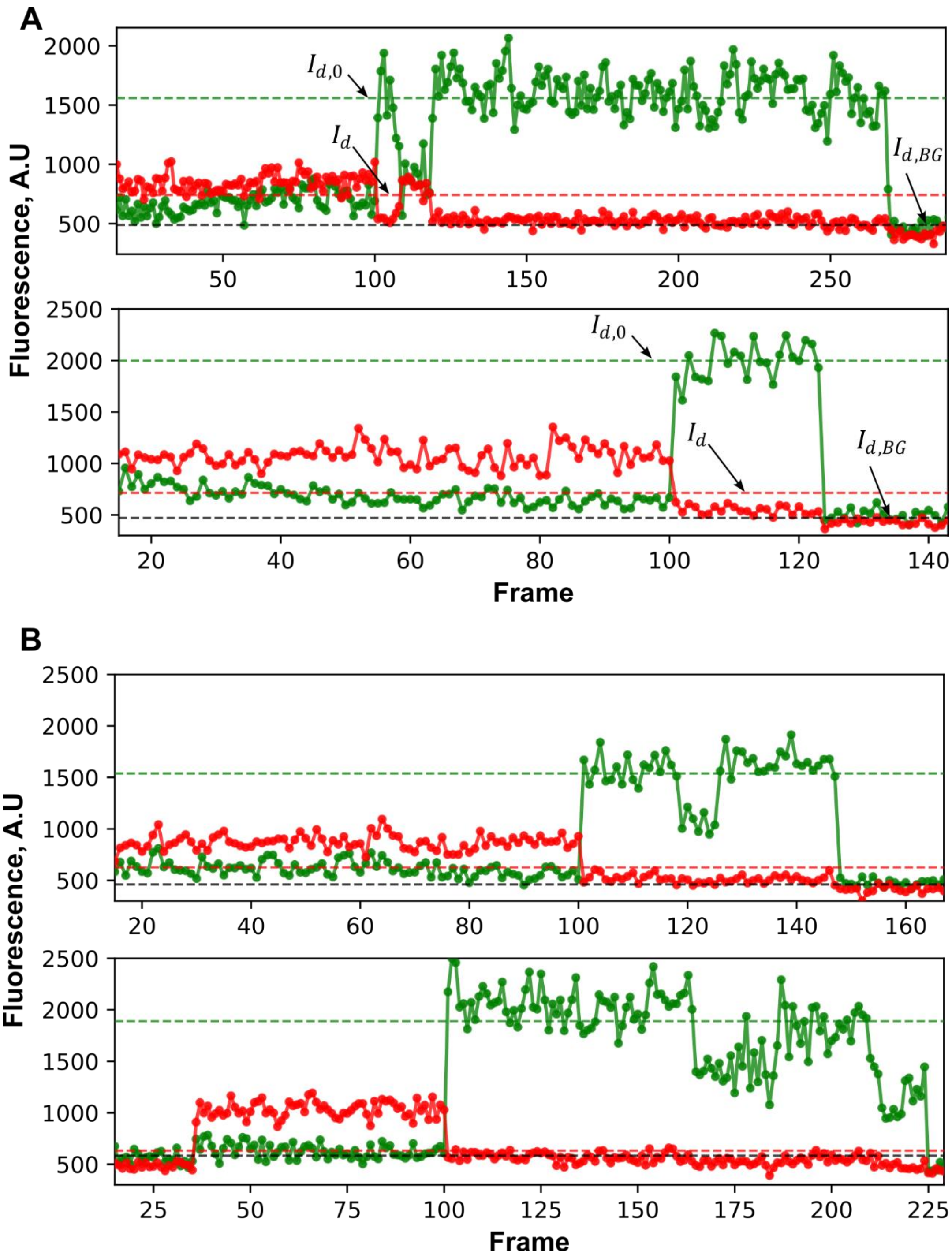


**Figure S4.** Selection of FRET traces by acceptor, donor photobleaching. (A) Example of traces used to calculate FRET efficiency showing single step increase of Cy3B channel with concomitant decrease in FRET channel average fluorescence intensity followed by single step decrease of Cy3B intensity upon photobleaching. Arrows indicate averaging of Cy3B intensity trace to derive values used to calculate FRET efficiency. (B) Examples of traces rejected from analysis due to Cy3B trace shifting between intensity values indicating dark states and intensities indicating more than one molecule.

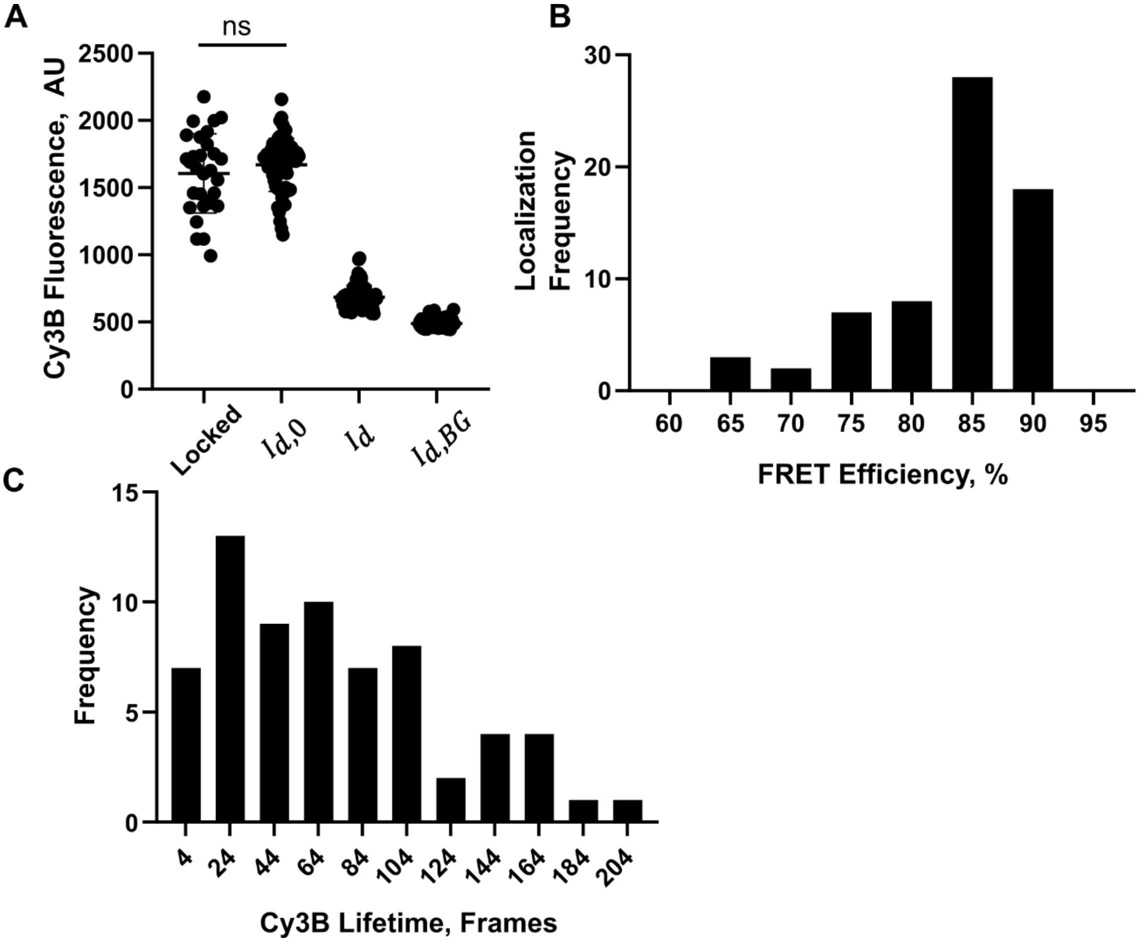


**Figure S5.** Analysis of single molecule intensity, FRET efficiency, and survival lifetime of LR probe Cy3B. (**A**) Fluorescence intensity of single molecule Cy3B traces upon locking strand induced opening of anchor strand hybridized to ligand strand with quencher(Locked), after single-anchor stranded acceptor photobleaching(Id0), during FRET(Id), and after Cy3B photobleaching(Id,BG). (**B**) Histogram of FRET efficiency calculated from single molecule Cy3B traces of single anchor strand. The FRET efficiency $\boldsymbol{E} = \frac{\boldsymbol{I_{d,0}} - \boldsymbol{I_d}}{\boldsymbol{I_{d,0}} - \boldsymbol{I_{d,BG}}}$ was calculated where $\boldsymbol{I_{d,0}}$ is the donor intensity after acceptor photobleaching, $\boldsymbol{I_d}$is the donor intensity during FRET, $\boldsymbol{I_{d,BG}}$ is the donor channel background intensity determined after donor photobleaching. (C) Cy3B exposure lifetime after as calculated by the number of frames where Cy3B is bright after A647N acceptor photobleaching.

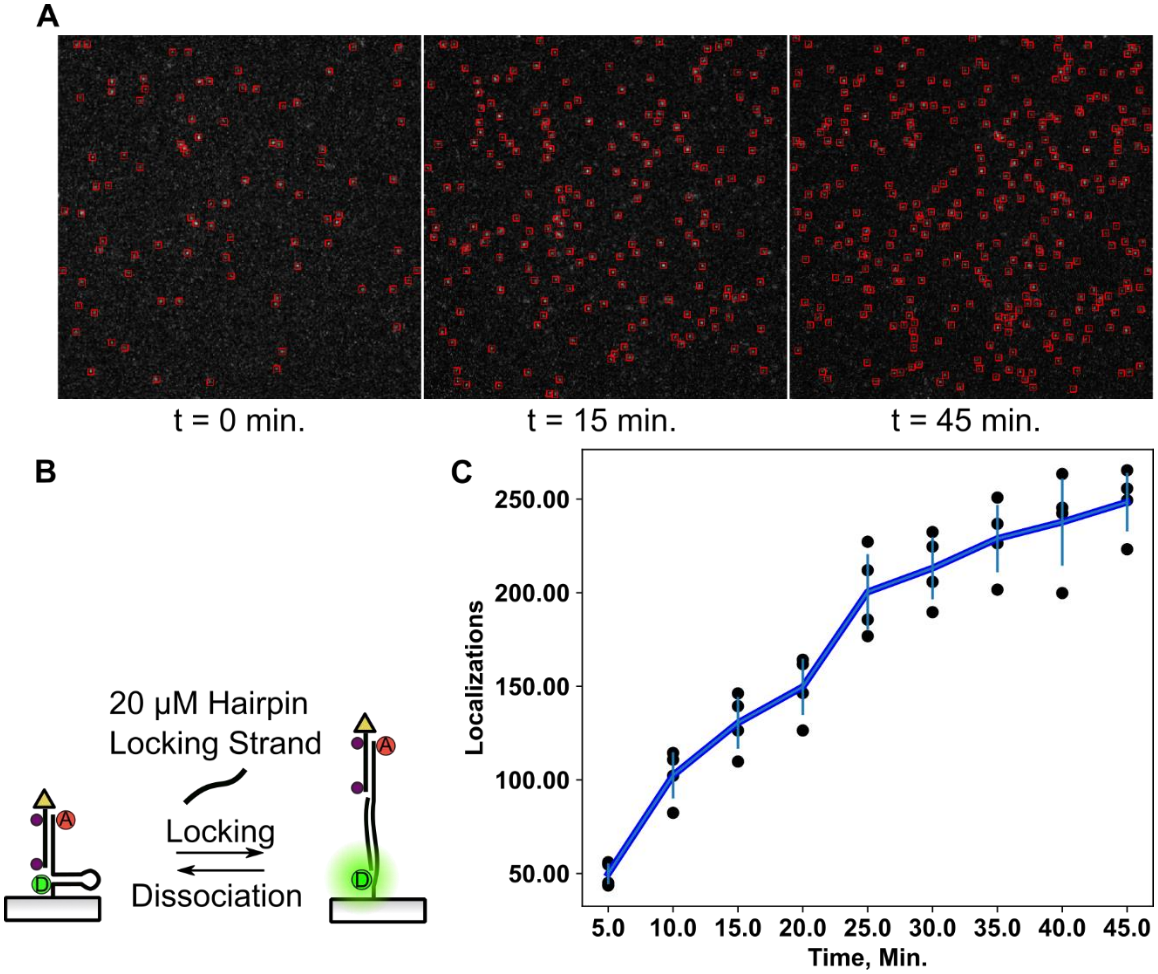


**Figure S6. LR probe surfaces exposed to hairpin locking strand.**
(**A**) Fluorescence micrographs of LR probe surfaces exposed to 20 µM 17 bp locking strand, added at t = 0, complimentary to the hairpin sequence (**Table S1**) with Picasso localizations boxed in red at different time points. (**B**) Diagram of locking strand induced hairpin opening. (**C**) Localizations over time for 4 surfaces(black dots) with the average(dark blue line) and standard deviation(light blue horizontal line).

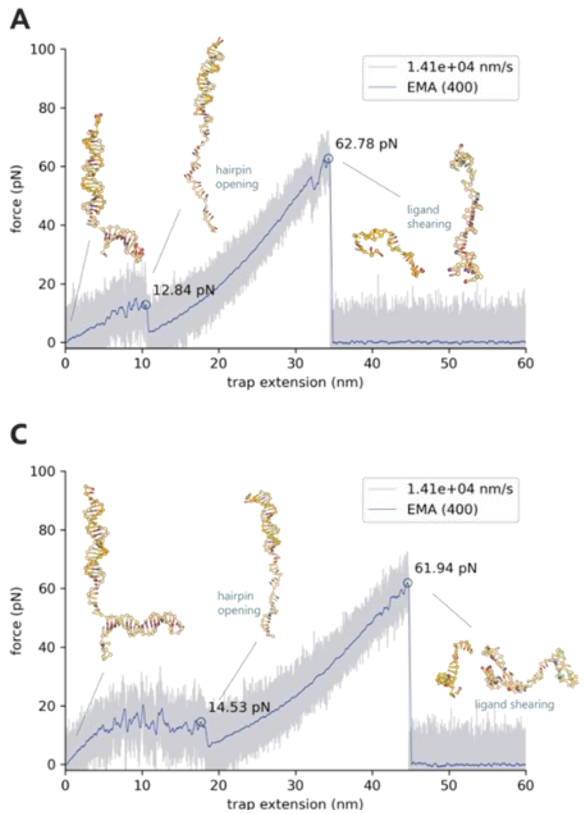


**Figure S7.** Force-extension curves of the LR probe as modeled by oxDNA. (A,B,C) Force-extension curve of LR probes with indicated force loading rate at each terminus. Gray: Raw data, Blue: Exponential moving average of 400 data points. Each graph represents the three separate simulation runs.

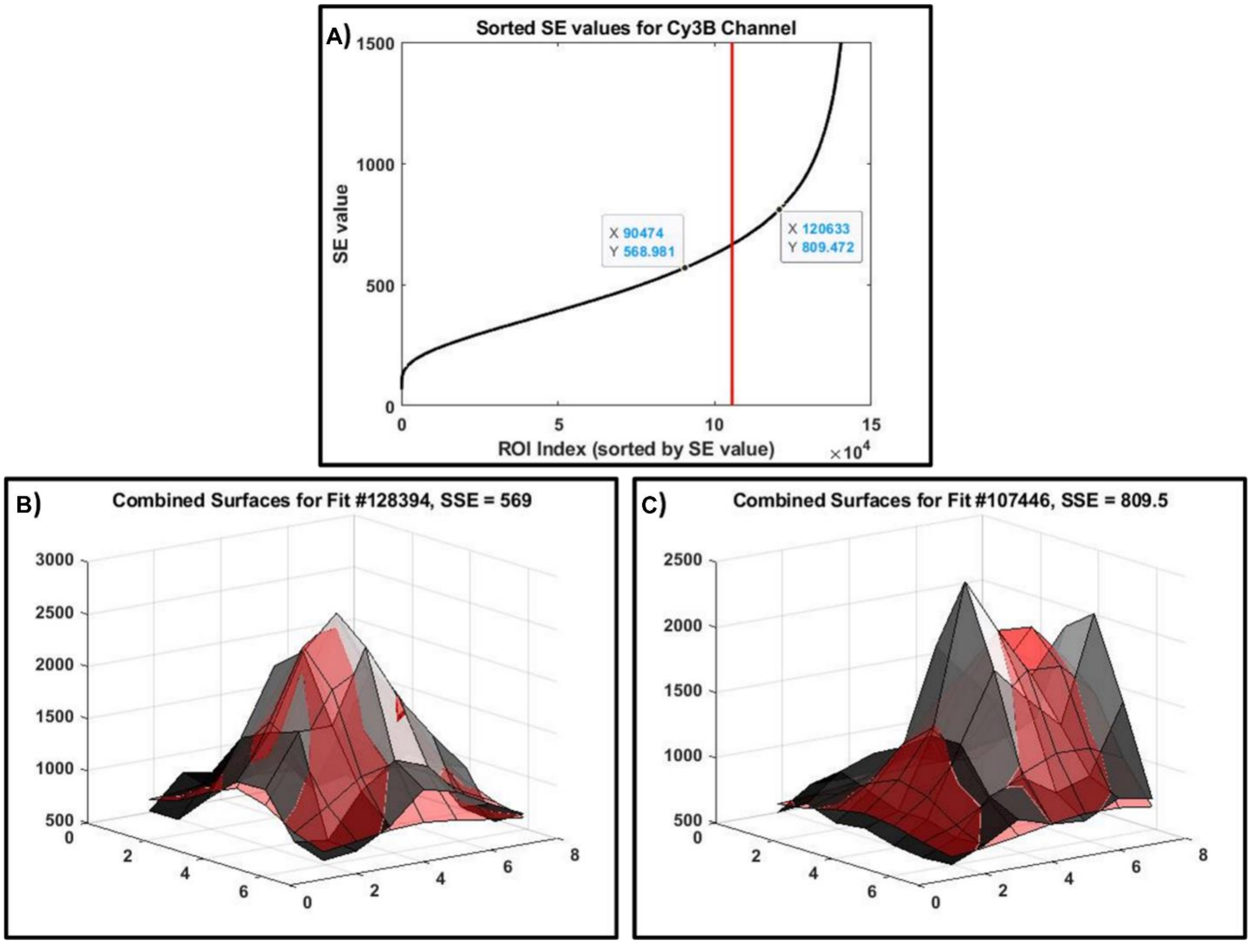


**Figure S8.** Identification of clean, single molecule events from gaussian fit data for further single molecule analysis. (A) Sorted SE values for one measurement's worth of ROIs in the Cy3B channel. The vertical red line depicts the 70th percentile threshold. Marked points have their surfaces and fits plotted in Figures S8B and C. (B) Surface and fit for an ROI matching the 60th percentile of SE values, this data was used in further single-molecule analysis. (C) Surface and fit for an ROI matching the 80th percentile of SE values, this data was excluded from further single-molecule analysis.

**SI note 1 – LR Probe Concept**

Our force loading rate (LR) probe is designed using two oligonucleotides that share a complementary domain (**Figure 1A, Table S1**). The top oligonucleotide (ligand strand) is chemically modified with a terminal cyclic peptide (cRGD) peptide to mediate integrin binding, while the bottom strand (anchor) covalently attaches the construct to the surface. The LR probe has three primary states which correspond to different force thresholds and can be identified by unique fluorescent signals: Closed, Open, and Sheared (**Figure 1C-E**). When the LR probe is at mechanical rest ($F$ <4.7pN) the signal from both fluorophores is quenched by the corresponding BHQ2 dye (**Figure 1C** – closed). Upon integrin-ligand complex formation and a small ($F$ >4.7 pN) force application, the hairpin domain is unfolded resulting in a signal from the Cy3B donor fluorophore (**Figure 1D** – opened). Once this force has increased to a high enough force threshold ($F$ >56 pN) the ligand strand is sheared and the hairpin domain refolds producing a FRET signal from the donor-acceptor Cy3B-Atto647N pair (**Figure 1E** – sheared).

The key functional feature of this LR probe is that it generates an initial Cy3B turn-on signal at low magnitude of $F$ >4.7 pN (opening event), and then this signal is dampened and followed by a FRET response when the force exceeds 56 pN (shearing event). Importantly, the appearance of both these signal events come from the same integrin-ligand interaction, and thus the force loading rate for each individual interaction can be inferred by taking the time difference between the two events. This process produces a sequence of fluorescence signals illustrated in the trace in **Figure 1F** which is used to calculate the force loading rate.

Table S2. Sequences of modified oligonucleotids as custom synthesized by Integrated DNA Technologies.

**<u>These files are being withheld by the authors until after peer-reviewed publication.</u>**

**SI note 2 – LR Probes Report Integrin-Ligand Tension**

**These files are being withheld by the authors until after peer-reviewed publication.**

**Figure S9. Loading rate probes report cell tension in Cy3B, A647N (640 nm excitation), and sensitized FRET fluorescenace channels.**

<u>**These files are being withheld by the authors until after peer-reviewed publication.**</u>

**Figure S10. LR probes report ligand specific opening and shearing at low probe density.**

<u>**These files are being withheld by the authors until after peer-reviewed publication.**</u>

**Figure S11. Localizations of different LR probe anchor constructs.**

<u>**These files are being withheld by the authors until after peer-reviewed publication.**</u>

**SI note 3 – Automated Analysis of Single-Molecule Data**

Custom Matlab codes were written and implemented on each measurement for automated calculation of the single-molecule loading rate. Input parameters for the codes were first optimized on a representative dataset and then applied uniformly throughout all datasets during analysis. The custom code was written to perform 4 primary steps:

1) Finding optimal single-molecule events using Gaussian fitting
2) Obtaining time-traces of single-molecule events
3) Identifying key timepoints within time-traces and sorting events
4) Compiling kinetics of LR probe-rupturing event

The analysis used to perform these four steps and the logic behind them are described in detail below:

Note: For the following SI sections each step of the analysis will be performed on a single representative data set for the sake of simplification. The shown data analysis process is repeated in an identical manner across all data within the experiment.

#### 3.1 Analysis of SM Data 1: Finding optimal single-molecule events using Gaussian fitting

The first step is to identify single molecules which can be analyzed independently from all other molecules. Fluorescent spots were identified by scanning the image for localizations which satisfied three criteria: 1) Intensity above a certain threshold (1.85 or 2.4 times the value of the background for the FRET and Cy3B channels, respectively), 2) local maxima in the x and y-direction, and 3) a peak prominence above 20 counts (to minimize the effects of shot noise). Pixels which satisfied these criteria were marked as a region of interest (ROI), and a 7x7 region centered on the localization was fit to determine the quality of that ROI. In general, a diffraction limited spot made from a fluorescent single molecule should be well fit with a 2D-Gaussian fit with the form (**Equation S1**):

Equation S1. 2D-Gaussian fit for a diffraction limited spot.

$$\boldsymbol{Fit(x,y)} = \boldsymbol{I} * \exp\left(-\left(\frac{(x-\boldsymbol{x_0})^2}{2(\boldsymbol{\sigma_x})^2} + \frac{(y-\boldsymbol{y_0})^2}{2(\boldsymbol{\sigma_y})^2}\right)\right) + \boldsymbol{b}$$

Where $x_0$ and $y_0$ are the Gaussian's center in x and y, respectively, $\sigma_x$ and $\sigma_y$ are the Gaussian spread in x and y, I is a normalization factor, and b is the background. In the above equation the variables of the fit are marked in red and adjusting these variables changes the output 2D Gaussian surface, F(x,y), which is then compared to the raw data in order to quantitatively determine a "best fit" (typically a local maximum within this parameter space). A representative ROI showing such a fit and how the individual parameters affect the fit is displayed below in **Figure S12**:

The "best fit" was quantitatively determined using a form of Maximum Likelihood Estimation (MLE) which was chosen based off of the assumption that noise in the measurement follows a Poisson Distribution[19], and follows the form:

Equation S2. Summed error from the Poisson distribution of the data and fit.

$$\mathbf{SE} = \sum\left(\boldsymbol{Data(x,y)} * \ln\frac{\boldsymbol{Data(x,y)}}{\boldsymbol{Fit(x,y)}}\right)$$

Where SE if the summed error (inverse fit quality), ln is the natural logarithm, Data(x,y) is the raw data, and Fit(x,y) is the fitted surface from **Equation S1**. The error is calculated at each pixel within the ROI (here a 7x7, so 47 total points) and those errors are summed to give FQ. Intuitively, the natural logarithm of the ratio Data(x,y)/Fit(x,y) changes sign depending on which is larger, and so Fit(x,y) is normalized to the total intensity of Data(x,y) to avoid fit convergence on extremely large/small values. Lastly, custom bounds were added to the definition of FQ to restrain its parameter space to physically meaningful values given the single-molecule nature of the measurement (e.g. similar $\sigma_x$ and $\sigma_y$ values, peaks within 1.5 pixels of the ROI center, S:N ratio > 1, etc.).

In a given measurement there are often upwards of a hundred thousand ROIs. Once each ROI is fit and assigned a summed error, the top 70$^{th}$ percentile of SSEs are selected for further analysis. **Figure S8** shows all SE values in the Cy3B channel for a representative experiment, along with the 70$^{th}$ percentile threshold

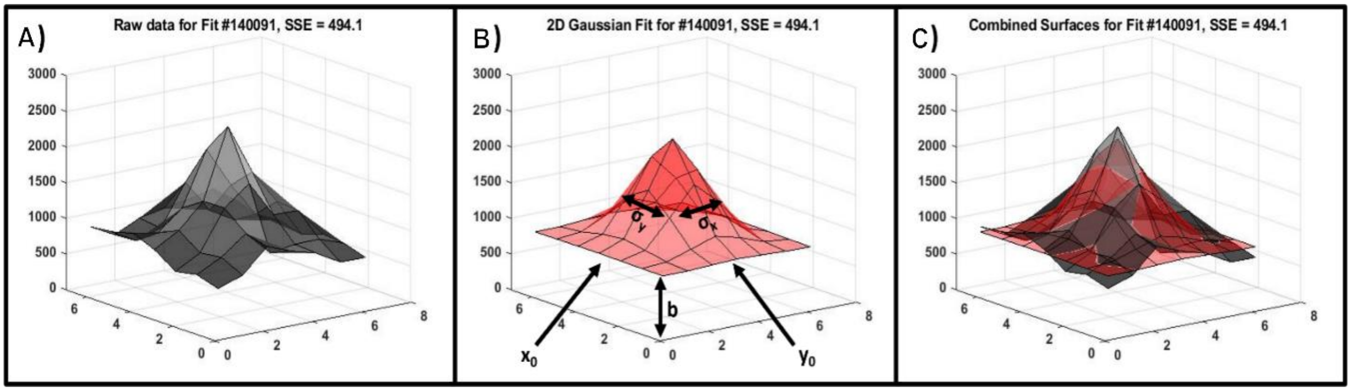


**Figure S12.** Single molecule fluorescence intensity gaussian fitting. (A) Raw data for a typical single-molecule spot captured in a single image. The shown molecule is the median SSE value of all fits, meaning it is a good representative selection. The area to be fit is a 7x7 pixel image. (B) Fit for the raw data using equation S 2, where the variables which define the fitted surface are depicted. (C) Combined raw data and fitted surface.

and representative fits which were slightly better and slightly worse than the threshold cutoff. As can be seen from **Figure S8 B**, fits below the 70$^{th}$ percentile of SE values fit well to the 2D Gaussian model for an expected single-molecule measurement. In contrast, the primary contributor to high SE values is attempting to fit two peaks to a single Gaussian, as seen in **Figure S8 C**. These molecules being close to each other would bias future analysis and decrease fidelity of the measurement, and are thus filtered out using their SE values as a metric for exclusion.

#### 3.2 Analysis of SM Data 2: Obtaining time-traces of single-molecule events

The previous section identified optimal single-molecule events, but does not yet take into account the temporal nature of the experiment. Namely, if the same molecule is observed in 10 frames the previous analysis will identify it as 10 different ROIs when in reality all ROIs belong to the same molecule. We grouped individual ROIs into clusters of molecules using a weighted DBSCAN algorithm, where the weighting of the dimensions clustered molecules that were within 1.2 pixels in the x and y-spatial dimensions, and 5 frames in the temporal dimension (to account for rapid fluctuations in LR probe conformations, dye blinking, etc.). After clustering, the fits which did not successfully cluster were removed (~2.5%) as they represent either noise or molecules with fluorescent signal which only exist for a single time-frame (faster than our temporal resolution). Compiling all relative positions of successfully clustered molecules at the single-molecule level is shown in (**Figure S13 A,B**), where we selected a cluster composed of a single molecule existing across hundreds of frames and histogram the Gaussian peak's location in the x and y-dimensions. As can be seen, the Gaussian peak's position remains relatively stable across the registered time lapse, as expected. We then compared the position of the Gaussian peak's position (the *fit* center) with the pixel that defines its ROI center (the *raw data* center) across all molecules within a measurement (**Figure S13 C**), which shows the precision of our localization. Since the fluorescent LR probes are immobilized on a surface, the positional distributions are due to a combination of noise, camera drift, etc. and validate the 1.2 pixel parameter used in the DBSCAN clustering (all fits within a cluster are within 1.2 pixels of each other, even when comparing different clusters' relative positions with each other, (**Figure S13 C**).

After clustering fluorescence events into single-molecule localizations the positions of all events within a cluster were averaged together to find the most consistent central position of that molecule (e.g., averaging all points in **Figure S13 A** and **B** for that cluster). Finally, to create a timetrace for that cluster,

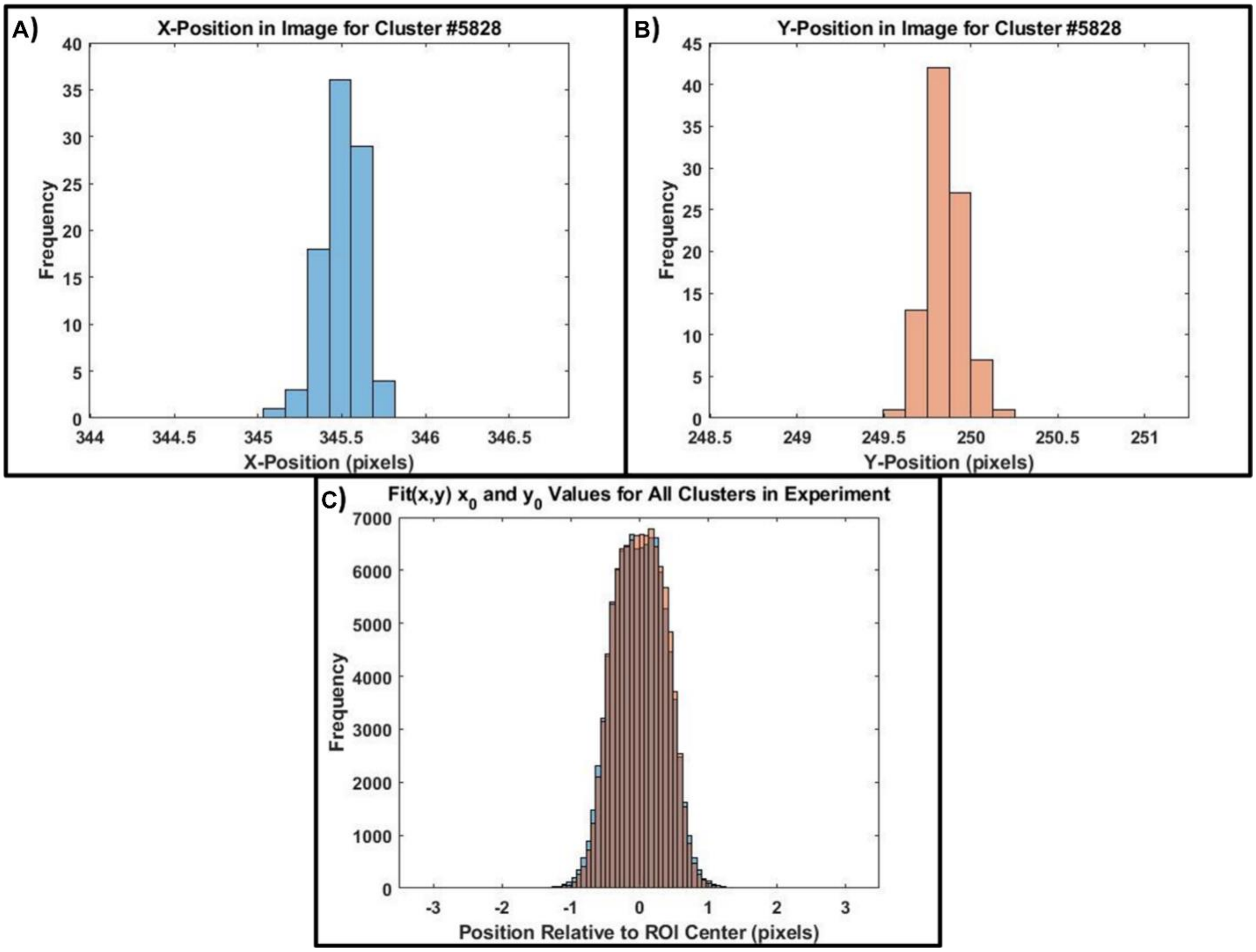


**Figure S13.** Distributions of the $x_0$ and $y_0$ values from fits using Equation S 2, where all values reported passed through the filtering process described. (A,B) The X and Y-Positions of a single cluster spanning multiple time frames (and therefore several fits) where all values are ascribed to an individual molecule. The x-axis shows the molecules absolute position on the images after registration. (C) The raw $x_0$ and $y_0$ positions for all fit molecules after filtering. The x-axis is relative to the center of the initial ROI obtained as described in this SI.

the intensities of the 3x3 area of pixels centered at the cluster's central position were summed for each frame across the timescale of the experiment to create an intensity time trace for that molecule (**Figure S14**).

The timetraces within **Figure S14** contains information on how each molecule is behaving during the timescale of the experiment. For instance, **Figure S14 B** (orange) shows a high intensity at the beginning of the experiment, suggesting that either the LR probe was opened *before* the image acquisition began (during the pre-incubation period while waiting for the cell to spread on the surface) or that the Cy3B dye for this probe began the experiment unquenched (due to the quencher photobleaching before data acquisition began or due to it not being labeled). In contrast, **Figure S14 C,D** show a probe where the Cy3B intensity starts low (quenched) and it becomes unquenched (most likely due to opening of the probe, but possibly due to infrequent photobleaching of the quencher). In all cases, the intensity goes up in a single step and decreases down to background in a single step, providing strong evidence that we are observing the LR probe dynamics at a single-molecule level.

Determining the timepoints when an intensity rises and falls, as well as the previously derived position of each single-molecule trace in both the donor and FRET channels is further analyzed in the next section, and the information is used determine kinetic information about the LR probe.

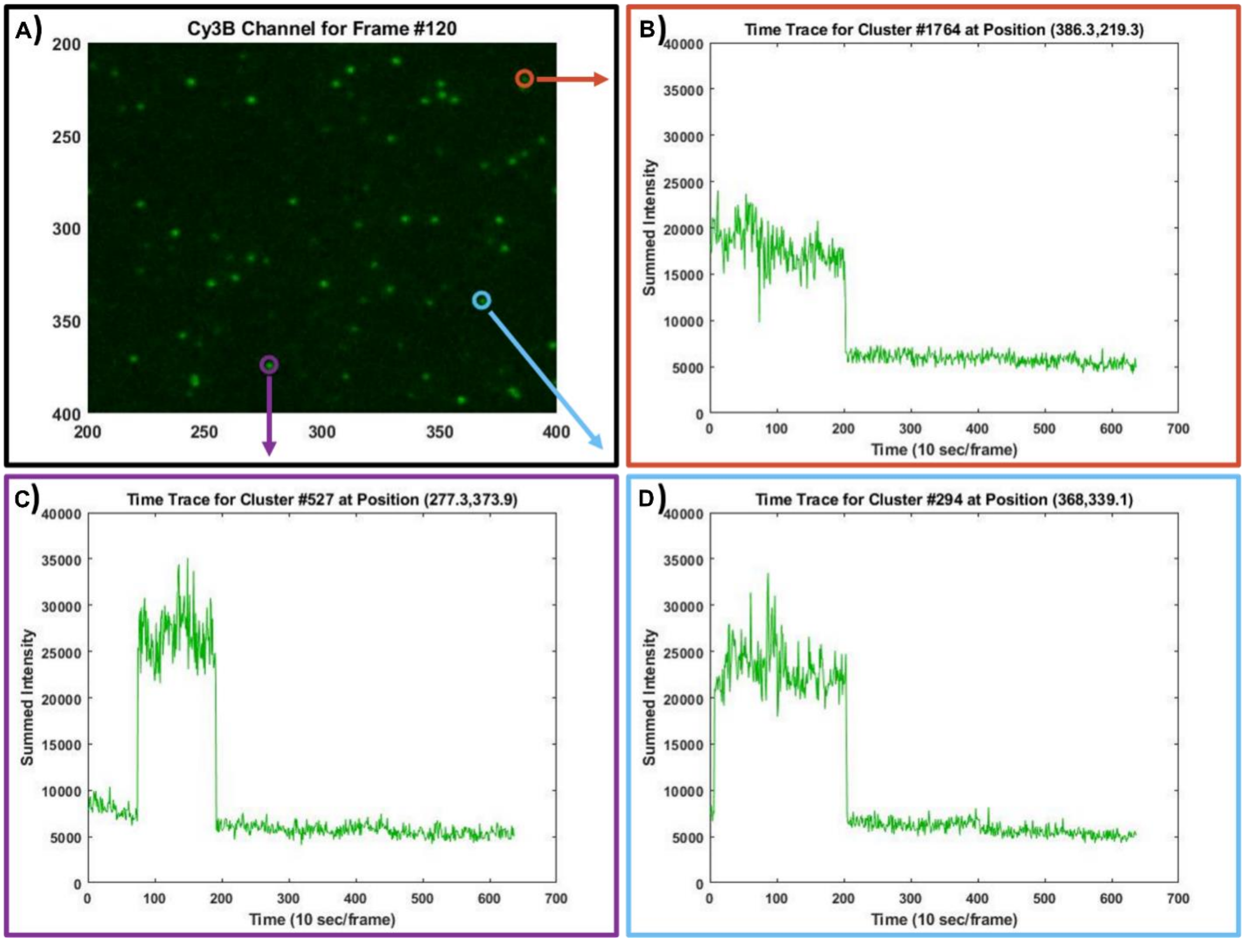


Figure S14. Time trace derivation from single-molecule events. (A) A 200x200 pixel zoom of the Cy3B channel at Frame 120 (20 minutes into the experiment). (B,C,D) Time traces for three randomly selected clusters, selected from the set of clusters centered within the 200x200 pixel range which were on during frame #120.

#### 3.3 Analysis of SM Data 3: Identifying key timepoints within time-traces and sorting events

In order to quantify when intensity shifts happened within a probe we employed a changepoint algorithm[20]. We performed this algorithm on all single-molecule events while setting the maximum

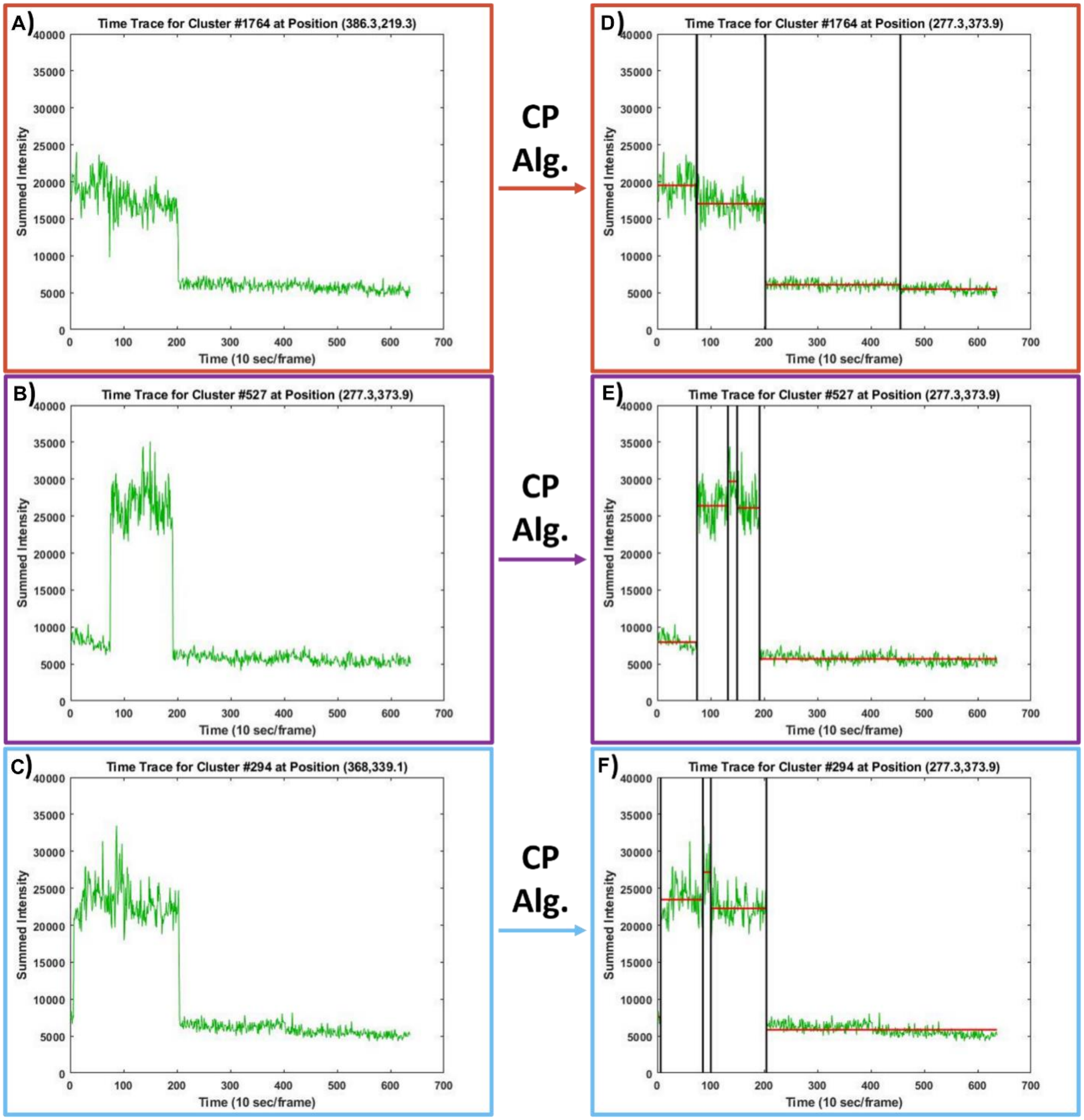


**Figure S15.** Single-molecule traces after implementing a change-point algorithm. (A,B,C) Initial single-molecule time traces reproduced from **Figure S14**. (D,E,F) Same traces after the change-point algorithm (CP Alg.) was applied. Black lines show the time points of each changepoint. Red lines show the average intensity value between changepoints.

number of intensity changes to 4 (off -> on -> off -> on -> off). This maximum number of changepoints was implemented to reduce the number of "false positives" due to the noise of single-molecule measurements, while still allowing the capture of all significant kinetic events within each single-molecule trace.

The same representative traces from **Figure S14** are reproduced in **Figure S15**, but with the before/after implementation of the changepoint algorithm (CP Alg.) visualized on top of the intensity traces. The timepoints for each changepoint are marked in black and, as can be seen, the CP algorithm has a tendency to "overfit" the intensity traces (**Figure S15 D,E,F**), which is why the constraint of a maximum of 4 changepoints was added. An example of this "overfitting" can be seen in **Figure S15 D**, where a changepoint is found at time frame ~450, where it detects a difference in background intensity. It should be noted that interesting fluorescent properties *may* be linked to these changepoints, and that these "overfit" changepoints are not necessarily false detections. However, our work is primarily interested in large intensity shifts resulting from probe opening/closing/rupturing events. To restrict our analysis to these events, it is beneficial to limit the maximum number of changepoints so only the largest intensity shifts are detected. In addition to finding the timepoints for each changepoint (black vertical lines), we also obtain the average intensity between changepoints (red horizontal lines). Taking the average intensity before/after each changepoint informs us on how significant the intensity shift is between detected changepoints, as well as the direction of those changepoints (on vs. off events).

#### 3.4 Analysis of SM Data 4: Compiling kinetics of LR probe-rupturing event

Having analyzed the Cy3B (donor) channel to filter single-molecule events, fit their super-resolved positions, create intensity timetraces, and obtain information on the intensity shifts as described above, the same analysis was repeated on the FRET channel to obtain two parallel data sets of the same dataset. A probe rupturing will follow the timeline of a probe opening (Cy3B turn-on) and the probe rupturing (Cy3B turn-off and FRET turn-on occurring simultaneously). We identified LR probe rupture events using the following criteria:

1) Two single-molecule cluster localizations in the Cy3B and FRET channels must be within 2 pixels of each other.

2) The Cy3B timetrace must experience a turn-off event.

3) The FRET timetrace must experience a turn-on event.

4) The turn-off/turn-on events in 2 and 3 must occur within a certain time of each other (threshold was within 2 frames).

For this representative dataset a total of 40 rupture events were found which satisfied the above criteria. 15 randomly selected unique traces are shown in **Figure S16** to represent the dataset. In these representative traces, the timepoints for probe opening (Cy3B turn-on) are shown in purple, while the average timepoints for criteria 2 and 3 (Cy3B turn-off, FRET turn-on) are shown in blue. The total amount of time the probe spent in the opened state before rupturing is the difference between the blue and purple timestamps (shown as ΔT in the figure titles in **Figure S16**), and has a time-resolution of 10 seconds (imaging framerate).

In addition to calculating the ΔT or probe-opening to probe-rupture, the time-traces also show additional evidence for single-molecule probes by analyzing the turn-off events. After rupturing, the probe has two dyes (donor and acceptor) which contribute to the FRET signal: if the donor photobleaches both signals should go to background (**Figure S16 C,D,G,K,L,M,O**) or if the acceptor photobleaches the donor signal should increase while FRET signal goes to background (**Figure S16 B,E,F,H,J,N**) which may or may not be followed by donor signal photobleaching, if this occurs on the timescale of the experiment. These step-wise transitions provides even further evidence that the timescales we are observing come from true single-probe measurements.

Lasty, the locations and timepoints of these probe-rupturing events can be mapped in real-time onto the cell's position (as in **SI Video 1, 4-10**). These videos shows an RICM image of the cell taken in parallel with the donor and FRET channels. The red circles overlapped on top of the RICM show the positions of the clusters, and the frame in which the red circles appear correspond to time of probe rupturing (blue line, **Figure S16**). The video also contains a real-time summation of the total number of probe-rupture events occurring within the measurement, as well as the frame count. As is visually apparent from the video, the probe-rupture events mostly occur within the region occupied by the periphery of the cell, where tension is expected to be strongest, providing further strong evidence that these events are real and not due to noise inherent to single-molecule experiments.

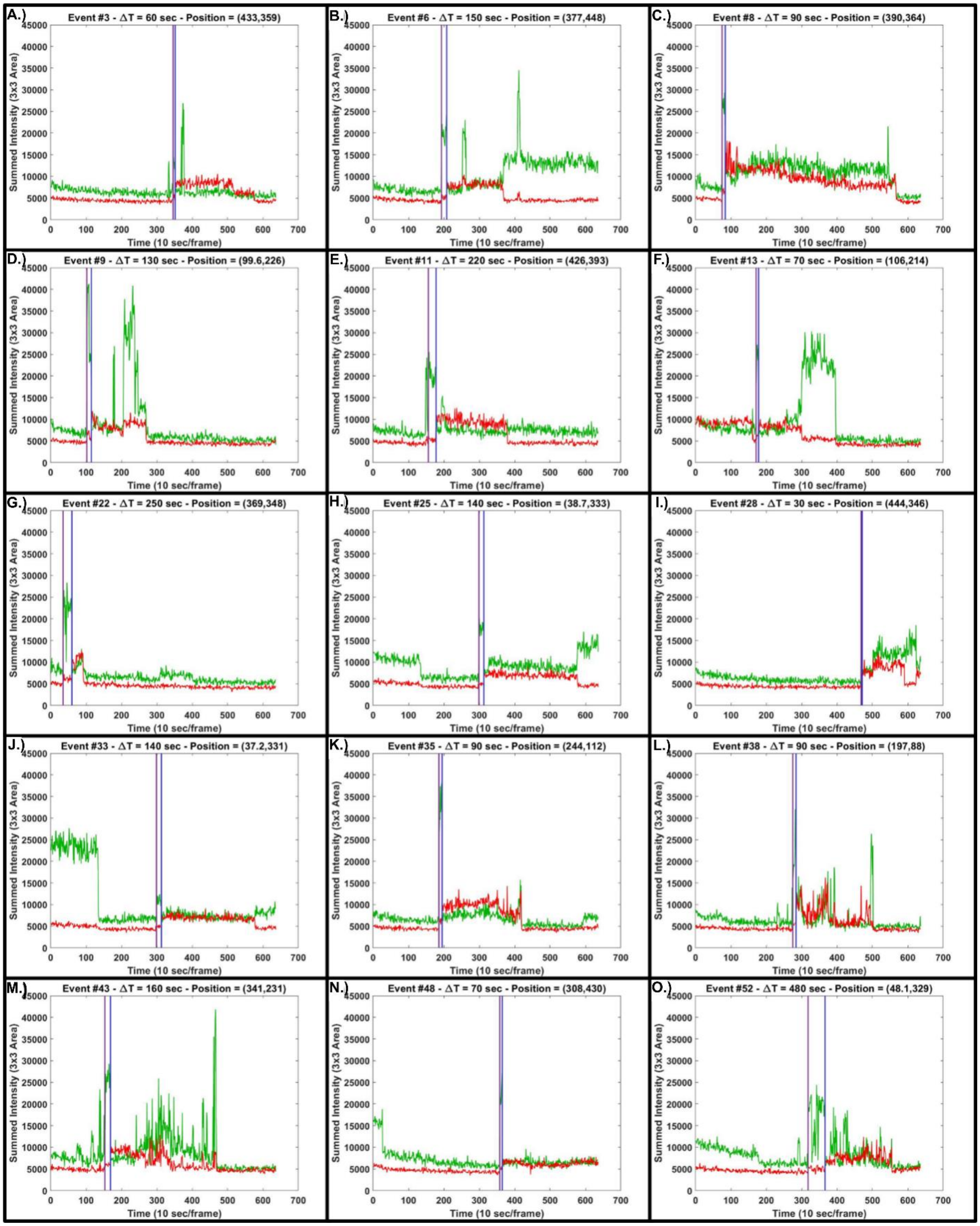


Figure S16. Timetraces for Cy3B (green) and FRET (red) channels for rupture events, as identified by the 4 criteria listed above. The timestamp for the probe opening (Cy3B turn-on, purple) and the probe rupturing (Cy3B turn-off and FRET turn-on, blue) are shown as vertical lines. The cluster event index, rupture time (ΔT = blue – purple), and cluster position are shown in image titles.

**3.5 Analysis of SM Data 5: Identifying Closed→Opened→Closed Transitions**

Further events can be extrapolated from the data by setting a different set of criteria from those outlined in Section 4: Compiling kinetics of LR probe-rupturing event. For instance, identifying the probes which undergo a closed→opened→closed transition were found using the following criteria:

1) The donor signal must start at a signal close to background (low-signal state, for our experiments this was 7,000 counts across the 3x3 pixel region).

2) The signal must then increase in intensity by at least double (high-signal state)

3) Finally, the signal must then return to the low-signal state.

Additional selection criteria were applied to filter the events to make sure the signal came from a single molecule, further criteria included:

1) Transition events must not be within 2 pixels of each other (to avoid misinterpretation of overlapped of signal from single-molecule LR probes).

2) Event location must remain within camera FOV throughout the experiment (to remove false turn-on or turn-off signals due to camera drift during experiment).

For other experiments, additional criteria may be added are sensible within those parameters.

Lastly, for obtaining these timepoints and intensities of these transitions a changepoint algorithm was used, with a high error threshold set to find the most obvious changepoints. As the experiments can span a large timescale (sometimes > 600 frames), and a high-signal state can exist for as short as 1 frame, approximately 10% of events were so short-lived as to be missed by the algorithm with broad parameters. To compensate for this, a “second-pass” was performed on events which did not identify a high-signal state, and individual intensity points were identified which exceeded twice the intensity of their assigned state. Such high-intensity points were assigned to the opened state, and the dataset of filtered closed→opened→closed transitions is represented in **Figure 4d**.

From **Figure 4d** and **SI Video 3**, it is clear that some events assigned closed→opened→closed transitions follow photobleaching kinetics (**Figure 4d**) and/or occur outside the area of the cell (**SI Video 3**). These events are likely due to photobleaching of the $Q_1$, which allows the donor to turn on, and results in a false "opened state". It is relatively simple to "background subtract" these photobleaching events, through the fits shown in **Figure 4d**: The relative area under each fit shows approximates the relative population for each event type (opened vs bleaching). Thus the final numerical values reported in the manuscript and represented in **Figure 4c** are the total number of events multiplied by the relative population of opened:bleach events (in our case, 77.4%).

**SI Method 1 – LR Probe Synthesis**

**Figure S17.** Synthesis of Cy3B-Biotin-Azide.

**Figure S18.** LR anchor HPLC chromatograms.

**Figure S19.** Synthesis and purification of cRGD-Azide.

**Figure S20.** LR ligand HPLC chromatograms

**Figure S21.** cRGD-dsDNA-biotin HPLC chromatograms.

**These files are being withheld by the authors until after peer-reviewed publication.**

**SI method 2 – LR Probe Surface Preparation**

**These files are being withheld by the authors until after peer-reviewed publication.**

**SI method 3 – OxDNA Modeling of the LR Probe**

In addition to using DNA secondary structures and sequences with previously studied force induced transitions[2, 10], we modeled the opening force and the ligand shearing force of our novel construct using the oxDNA2 model[11]. MD simulations were run on the annealed DNA hairpin strands (**Figure S7**) at an extension rate of 1.41 x$10^4$ $nms^{-1}$ along the z-axis.

Temperature and [Na+] were set to 37°C and 0.156 M respectively to mimic in vitro experimental conditions and the following parameters were used in oxDNA(**Table S2**). The terminal nucleotides with the ligands and anchor were added with harmonic traps each with stiffness constants $k_1$ and $k_2$ (equivalent to springs in oxDNA) of 11.40 $pNnm^{-1}$. The combined effective trap stiffness $k_{eff}$ can be calculated using **Equation S3,** where $k_1$ and $k_2$ are the stiffness constants of the two traps between which the LR probe construct is pulled.

**Table S2.** Settings used for oxDNA modeling of LR probe force transitions.

| | |
|---|---|
| Steps | 5e9 |
| diff_coeff | 2.5 |
| thermostat | john |
| T | 37C |
| interaction_type | DNA2 |
| Newtonian_steps | 103 |
| use_ave_seq | 1 |
| verlet_skin | 0.05 |
| salt_concentration | 0.156 |
| dt | 0.005 |

Equation S3. Combined trap stiffness

$$\frac{1}{k_{eff}} = \frac{1}{k_1} + \frac{1}{k_2}$$

An extension rate of $1.4\times10^4$ nm/s along the z-direction was used to move one of the traps on the nucleotide to which the cell adhesion ligand is attached. To obtain the force-extension curve in each simulation, we extracted the extensions of harmonic trap from the attached nucleotides and then projected it along the z-axis. The force is calculated by multiplying the total projected extension with k*eff*. The data points were then smoothed with a 400-point exponential moving average (EMA) of the data points using python (10.1038/s41586-020-2649-2). The rupture force was estimated by picking the peak at the point of rupture using SciPy find_peaks module (10.1038/s41592-019-0686-2). From the simulations, the opening of hairpins remained within the range of 12-16 pN (**Figure S7**). The difference in hairpin opening force with regards to the experimental estimates can be reasoned using the following factors. 1) Experiments use a constant force clamp to estimate the 50% probability of hairpin dissociating at a particular force within a defined interval of time (e.g., 2 sec) whereas in simulation the probes are far from equilibrium. This leads to higher force estimation for hairpin opening opening[12-13]. The constant force is incrementally increased in experiments, while the simulation experiments were performed by applying a defined velocity to the nucleotide. In a typical force ramp setup, the rupture force of a DNA duplex is highly dependent on the loading rate (i.e., the higher the loading rate the higher the rupture force). 2) The loading rates used in simulation are much faster compared to experiments due to computational resource limitations.[14-15] 3) oxDNA simulations have an inherent level of stochasticity that results in small variations in the resulting force of mechanical transitions(**Figure S7 A-C**). These simulations yield a force difference between the unfolding and shearing events of 47.69 ± 1.73 pN, near to the 51.3 pN force difference inferred from the previously reported mechanical transitions of 4.7 and 56 pN for the opening and shearing domains respectively.

### 4. ESI-MS

**<u>These files are being withheld by the authors until after peer-reviewed publication.</u>**

### 5. Live Cell Imaging

Cells were detached from plates 0.25% (wt/vol) trypsin, 2.21 mM EDTA (corning) and resuspended in normal culture media at a 1:20 v/v ratio, centrifuged at 250 X G for 4 minutes and resuspended in Fluorobrite DMEM (Gibco A1896701) supplemented with 5% serum imaging media. The imaging well was washed with 200 μL imaging media before adding 400-800 cells to the 384 well plate well (area density of 0.32-0.64 cells/cm2). Cells were then incubated for 25 min. at 37 °C 5% CO2 for 25 minutes before imaging.

Cells were then imaged at room temperature on a Nikon Eclipse Ti microscope driven by the NIS Elements Advanced Research software. The microscope objective used was CFI Apo 100X (numerical aperture 1.49) objective (Nikon). The optical system includes a total internal reflectance fluorescence (TIRF) variable mirror launcher and a Nikon Perfect Focus System, an interferometry- based device which corrects z-drift of the stage. Reflection interference contrast microscopy(RICM) images were captured using a filter cube(Chroma 97270 SRIC C168785) and by a Lumen Dynamics X-Cite 120 LED light source. Fluorescence excitation of samples through the optical system was carried out, 561nm (50 mW, used at 0.5% LP), and 640 nm (100mW, used at 1% LP) lasers. A Chroma Quad (Chroma TRF89901 ZT405/488/561/640rpc) TIRF filter cube was used in the imaging acquisition of all three fluorescence channels. The optical configuration included the Andor TuCAM dual camera system comprised of two Andor DU-897 X-9319 camera with a conversion gain of 3 and gain multiplier of 800 imaging at 50-100 ms exposure times. A Chroma ZT647rdc dichroic mirror was placed in the optical path to deflect light of wavelength < 640 nm, comprising the emission of Cy3B, to the second camera **(Figure S1**).

To correct for the offset between image planes of the two cameras, slides with Tetraspec Beads (Invitrogen T7281) fluorescent in both channels were imaged and planes were aligned using an affine transform based algorithm on the imageJ software[17]. Timelapse Images were drift corrected using an imageJ plugin based on software written by Preibisch *et. al.*[17]. Cells were imaged at 10, 20, and 60 second intervals between fluorescence acquisitions taking two frames of three fluorescence images at each acquisition interval: A647N, then Cy3B and Cy3B sensitized FRET to A647N simultaneously on both cameras. To ensure that the timepoint of the opening and shearing events are not due to stochastic intensity fluctuation two steps were taken. The first is that we employed a dual camera system which splits the emission of Cy3B and A647N to two separate cameras which record Cy3B and fluorescence and FRET micrographs simultaneously (**Figure S1**). This ensures that the Cy3B and FRET channel intensities are synchronized in time. The second is that for every frame which is separated in time by the frame interval, two Cy3B and FRET fluorescence micrographs were taken separated by a 100 ms exposure time delay and averaged to produce a micrograph with an average intensity between the first and second micrograph at the specific frame. We applied this measurement to cells spreading on LR probes at 10, 20, and 60 sec. frame intervals and found that 10 sec. was sufficient to capture Cy3B opening traces that lasted many frames. (**Figure S22-S25**).

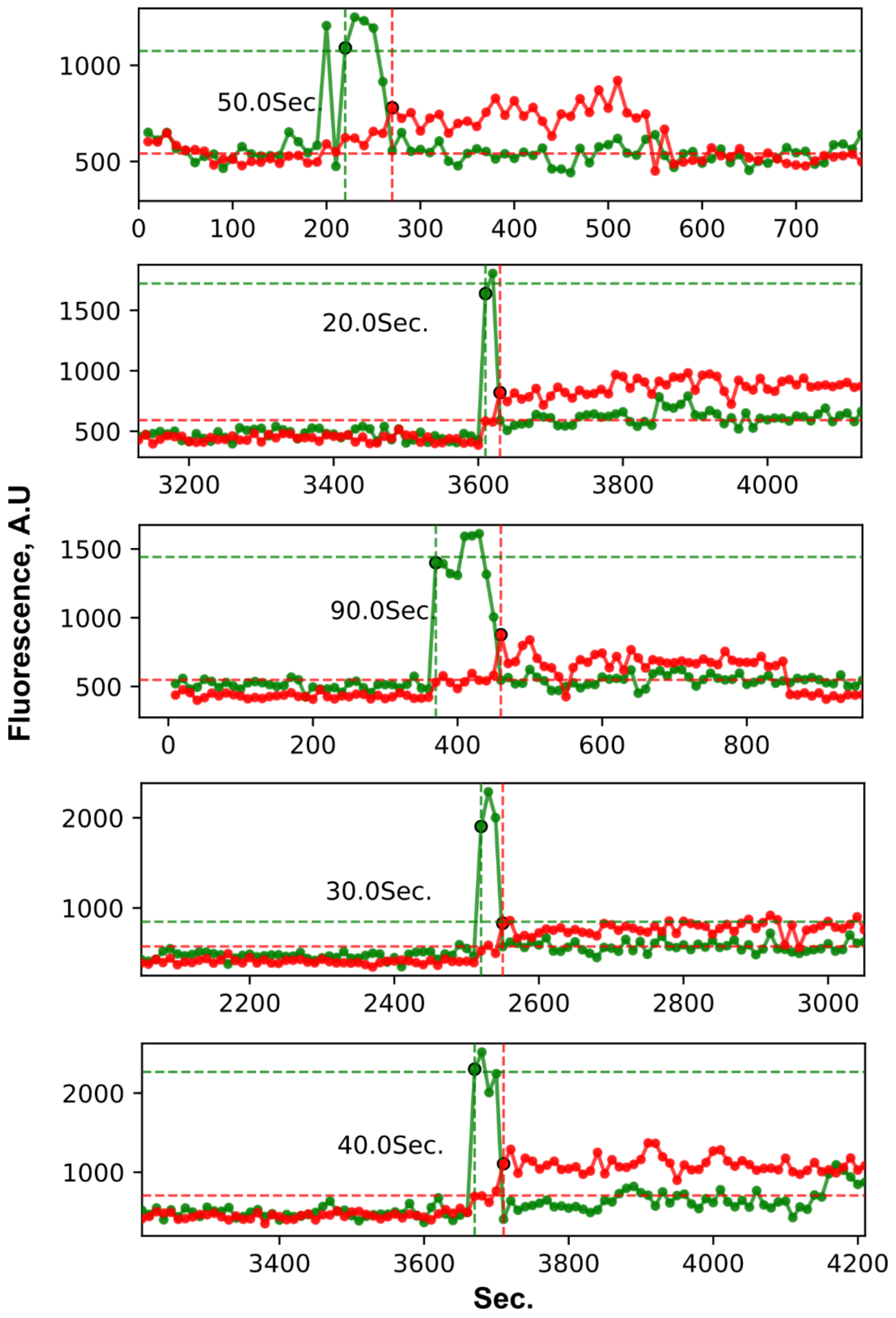


**Figure S22.** Representative loading rate trace localizations at a 10 sec. frame interval.

Loading rate trace localizations with the time between shearing and opening demarcated as calculated from the difference between the instance of Cy3B(green trace) turn on (green dashed vertical line) and the instance of simultaneous Cy3B decrease followed by FRET channel(red trace) increase(red dashed vertical line). Cy3B average intensity during turn on demarcated by vertical dashed line, and average intensity during FRET demarcated by red dashed line.

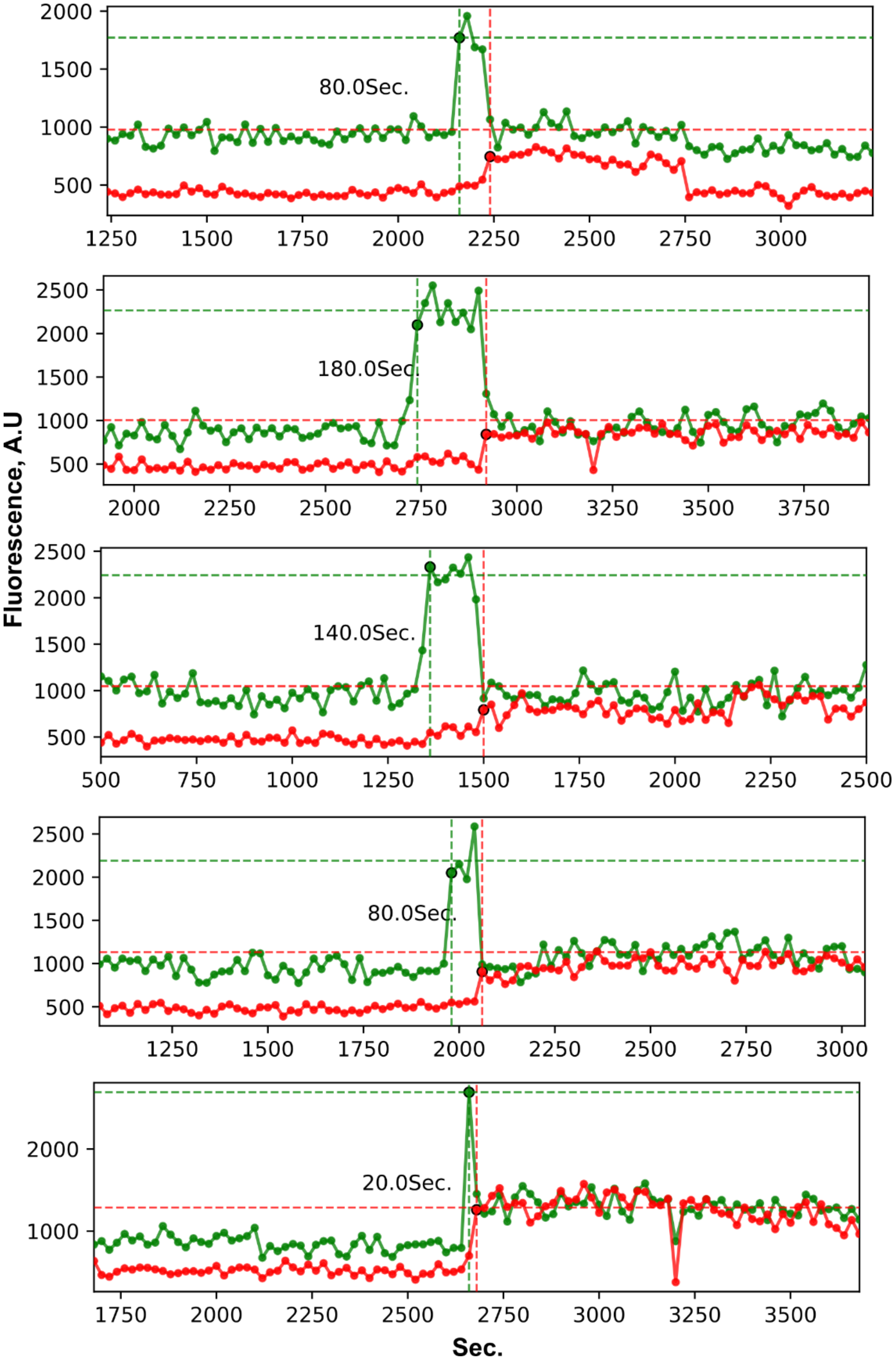


**Figure S23.** Representative loading rate trace localizations at a 20 sec. frame interval. Loading rate trace localizations with the time between shearing and opening demarcated as calculated from the difference between the instance of Cy3B(green trace) turn on (green dashed vertical line) and the instance of simultaneous Cy3B decrease followed by FRET channel(red trace) increase(red dashed vertical line). Cy3B average intensity during turn on demarcated by vertical dashed line, and average intensity during FRET demarcated by red dashed line.

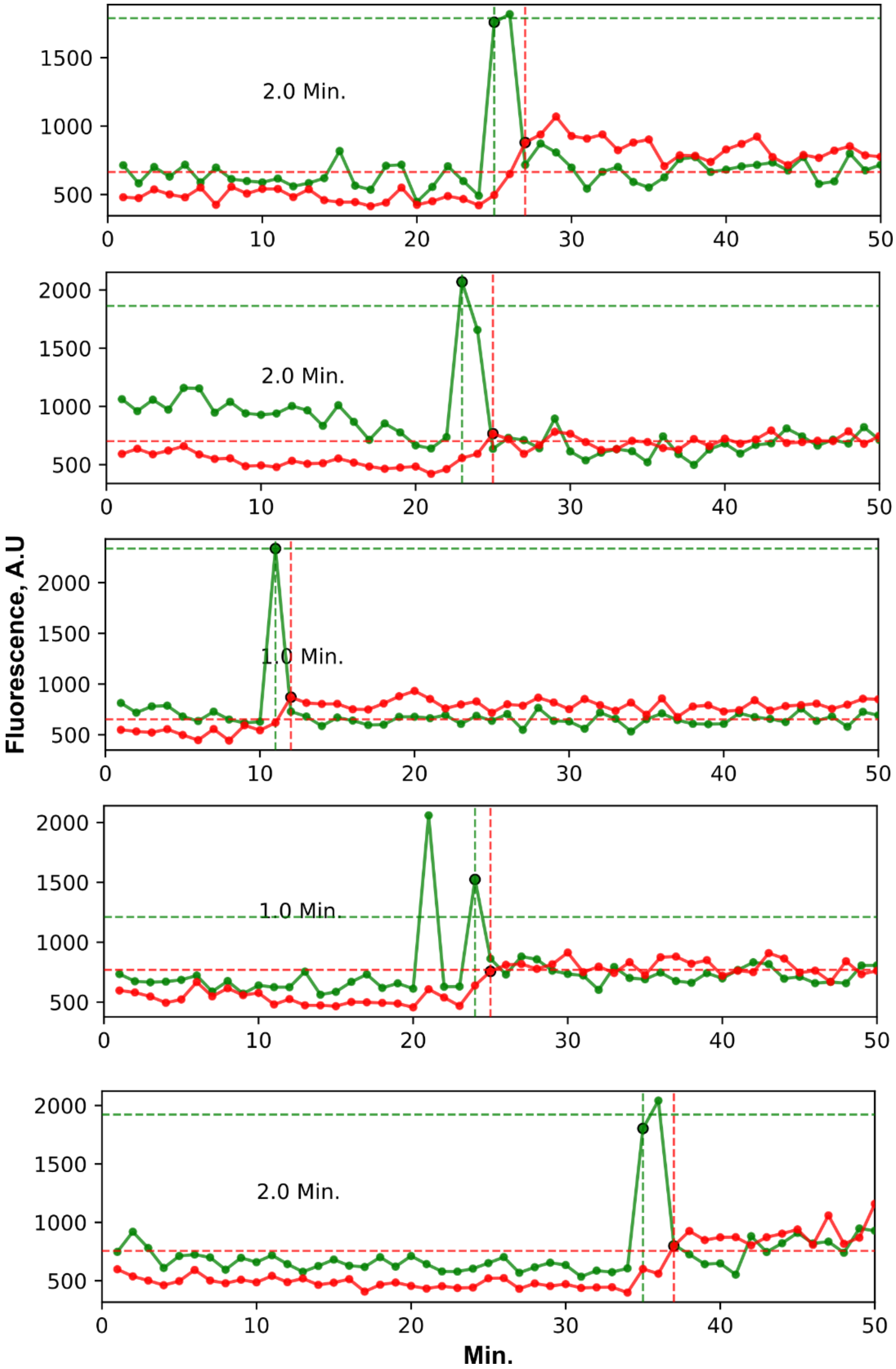


**Figure S24** Representative loading rate trace localization at a 1 min. frame interval. Loading rate trace localizations with the time between shearing and opening demarcated as calculated from the difference between the instance of Cy3B(green trace) turn on (green dashed vertical line) and the instance of simultaneous Cy3B decrease followed by FRET channel(red trace) increase(red dashed vertical line). Cy3B average intensity during turn on demarcated by vertical dashed line, and average intensity during FRET demarcated by red dashed line.

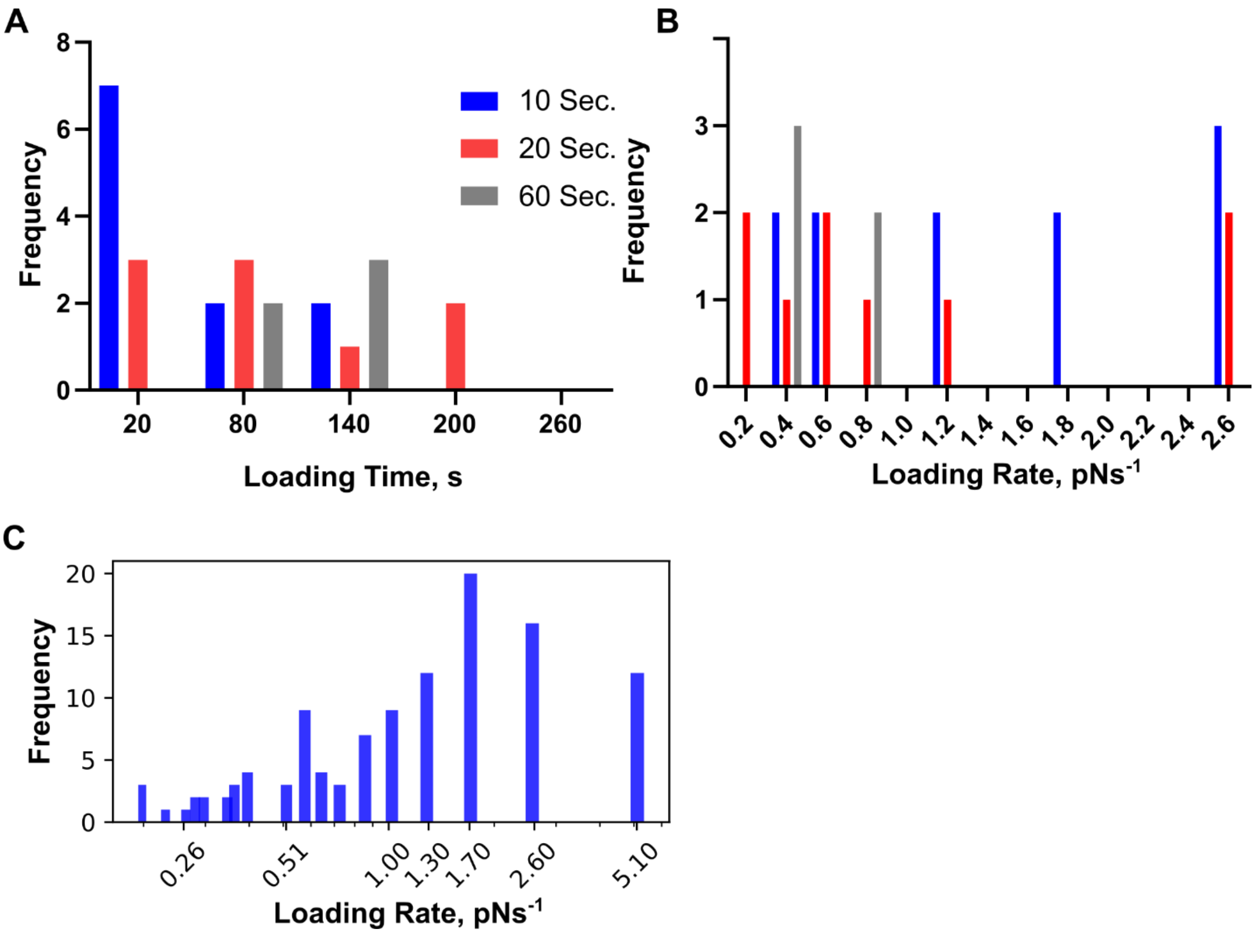


**Figure S25**. Manually identified Loading loading time and linear loading rate for different frame intervals. (A) Loading time as derived from opening-shearing traces for cells from 10, 20 and 60 sec, N =1 cell for each frame interval. frame intervals. (B) Loading rates assuming a linear force ramp between the opening and shearing signals and that opening occurs at 4.7 pN and shearing occurs at 56 pN. (C) Manually identified loading rate for cells collected at a 10 sec frame interval, N = 8 cells.

**6. Manual and Picasso assisted analysis of single-molecule data**

To qualitatively measure single-molecule fluorescence traces, a square 4X4 pixel (0.43X0.43 micron) square ROI was drawn around manually identified localizations. The average intensity of these localizations was measured over time to derive the intensity time traces. Only traces demonstrating single increase after acceptor photobleaching and single step decrease to background upon photo-bleach of Cy3B were used to calculate intensity values and FRET efficiency (**Figure S4**). Single molecule Cy3B brightness has been demonstrated to vary in fluorescence intensity of more than 20% for time traces on the order of minutes[1, 8]. The FRET efficiency $E = \frac{I_{d,0} - I_d}{I_{d,0} - I_{d,BG}}$ was calculated where $I_{d,0}$ is the donor intensity after acceptor photobleaching, $I_d$ is the donor intensity during FRET, $I_{d,BG}$ is the donor channel background intensity determined after donor photobleaching(**Figure S4 A**) [18]. The number of consecutive frames where Cy3B maintains high fluorescence after A647N acceptor photobleaching ($I_{d,0}$) was used to determine the average number of frames before Cy3B photobleaching, yielding an average of 71.1 frames, a standard deviation of 51.0 frames, and a standard error of the mean of 6.274 (**Figure S5 C**).
